# Long-read whole genome sequencing and comparative analysis of six strains of the human pathogen *Orientia tsutsugamushi*

**DOI:** 10.1101/280958

**Authors:** Elizabeth M. Batty, Suwittra Chaemchuen, Stuart D. Blacksell, Daniel Paris, Rory Bowden, Caroline Chan, Ramkumar Lachumanan, Nicholas Day, Peter Donnelly, Swaine L. Chen, Jeanne Salje

## Abstract

**Background:** *Orientia tsutsugamushi* is a clinically important but neglected obligate intracellular bacterial pathogen of the Rickettsiaceae family that causes the potentially life-threatening human disease scrub typhus. In contrast to the genome reduction seen in many obligate intracellular bacteria, early genetic studies of *Orientia* have revealed one of the most repetitive bacterial genomes sequenced to date. The dramatic expansion of mobile elements has hampered efforts to generate complete genome sequences using short read sequencing methodologies, and consequently there have been few studies of the comparative genomics of this neglected species.

**Results:** We report new high-quality genomes of *Orientia tsutsugamushi,* generated using PacBio single molecule long read sequencing, for six strains: Karp, Kato, Gilliam, TA686, UT76 and UT176. In comparative genomics analyses of these strains together with existing reference genomes from Ikeda and Boryong strains, we identify a relatively small core genome of 657 genes, grouped into core gene ‘islands’ and separated by repeat regions, and use the core genes to infer the first whole-genome phylogeny of *Orientia*.

**Conclusions:** Complete assemblies of multiple Orientia genomes verify initial suggestions that these are remarkable organisms. They have large genomes with widespread amplification of repeat elements and massive chromosomal rearrangements between strains. At the gene level, Orientia has a relatively small set of universally conserved genes, similar to other obligate intracellular bacteria, and the relative expansion in genome size can be accounted for by gene duplication and repeat amplification. Our study demonstrates the utility of long read sequencing to investigate complex bacterial genomes and characterise genomic variation.

## Introduction

### Background

*Orientia tsutsugamushi* is an obligate intracellular bacterial pathogen of the order Rickettsiales, family Rickettsiaceae which causes the life-threatening human disease scrub typhus. *Orientia* is transmitted by *Leptotrombidium* mites that occasionally feed on humans during the larval stage of development (“chiggers”), inoculate bacteria into the skin, and initiate infection. *Orientia* is maintained in mite populations by transovarial transmission. The mites normally feed only once on a vertebrate host, and cannot transmit bacteria directly from one host to another (Coleman et al., 2003). Bacteria propagate within endothelial cells, dendritic cells and monocytes at the site of inoculation, sometimes resulting in a visible red skin feature called an eschar (Paris et al., 2012). Bacteria subsequently spread through the endothelial and lymphatic system to cause a systemic infection characterised by lymphadenopathy, headache, fever, rash and myalgia, which typically begin 7-10 days after inoculation. The non-specificity of these symptoms makes scrub typhus difficult to diagnose based purely on clinical observations, and this is an important reason why the prevalence of scrub typhus has been historically under-recognised. Scrub typhus has now been shown to be a leading cause of severe fever and sepsis in studies in Thailand, India, China, Laos and Myanmar (Luce-Fedrow et al., 2018) and untreated or severe cases are associated with CNS infection, morbidity and death (Bonell et al., 2017; Dittrich et al., 2015). Locally acquired cases of scrub typhus have been reported in South America and the Middle East(Izzard et al., 2010; Weitzel et al., 2016), suggesting that the global burden of this disease may stretch beyond the traditionally known endemic areas of Asia and Northern Australia (Luce-Fedrow et al., 2018).

*Orientia tsutsugamushi* (previously *Rickettsia tsutsugamushi),* is distinct from other members of the Rickettsiaceae. The genus Orientia currently includes two known species, *O. tsutsugamushi* and *O. chuto,* the latter represented to date by a single strain isolated from a patient with a febrile illness contracted in Dubai (Izzard et al., 2010). High antigenic diversity among strains of *Orientia tsutsugamushi* is reflected in the poor immunological protection that recovered patients exhibit towards strains different from their original infection and, combined with a complex immune response that involves both humoral and cell-mediated immunity, this has hampered efforts towards vaccine development.

Despite its importance as a pathogen, few genomic analyses of *O. tsutsugamushi* have been published. The first whole genome sequence, Boryong, (Cho et al., 2007) reported a proliferation of type IV secretion systems in a repeat-dense genome of which 37.1% comprised identical repeats. A comparison of Boryong and the second complete genome, Ikeda (Nakayama et al., 2008), revealed similar repeats present in each genome, dominated by an integrative element named the *Orientia tsutsugamushi* amplified genetic element (OTage), and identified a core genome of 520 genes shared between the two *O. tsutsugamushi* strains and the 5 available sequences of other *Rickettsia* (Nakayama et al., 2010). Extensive genomic reshuffling was thought to have been mediated by amplification of repetitive sequences.

In comparison to other *Rickettsiae*, many of which have small and extremely stable genomes, *Orientia tsutsugamushi* has a large genome with an extraordinary proliferation of repeat sequences and conjugative elements. Some of the conjugative elements present in multiple copies across the genome are homologues of a gene cluster found in a single copy in *Rickettsia bellii*. Many of the genes in these elements are fragmented, suggesting they are non-functional (Darby et al., 2007). Other intracellular pathogens also contain repetitive elements, such as the mobile genetic elements in *Wolbachia* (Wu et al., 2004) and the tandem intergenic repeats in *Ehrlichia ruminantum* (Frutos et al., 2006). These mechanisms may evolve to increase genetic variability and aid immune evasion in bacteria which cannot easily take up novel DNA.

Larger collections of *O. tsutsugamushi* strains have been extensively studied using MLST and sequence typing of the *groES* and *groEL* (Arai et al., 2013) genes, and the outer membrane proteins 47kDa (also called HtrA or TSA47) (Jiang et al., 2013) and 56kDa (also called OmpA or TSA56) (Lu et al., 2010) genes. The 56kDa and 47kDa genes are highly immunogenic in human patients and animal models and have long been investigated as candidates for vaccine design, but high levels of diversity between strains, especially in the 56kDa gene, have limited the potential of developing a universal vaccine based on these epitopes.

Multiple studies in South East Asia have looked at the diversity of strains by MLST and 56kDa typing, and shown a high level of diversity, with many MLSTs unique to an individual strain (Duong et al., 2013; Phetsouvanh et al., 2015; Sonthayanon et al., 2010; Wongprompitak et al., 2015). Work in Thailand and Laos has shown recombination between MLSTs, as well as evidence for multiple infections in individual patients, implying that multiple strains may co-exist in mites (Sonthayanon et al., 2010). Comparisons of *56kDa* typing with MLST (Sonthayanon et al., 2010) and *47kDa* (Jiang et al., 2013) also show low congruence between methods, suggesting that single gene typing of *Orientia* may not represent the true relationships between strains; by extension, a 7-gene MLST scheme may not capture the full set of genomic relationships among strains.

Attempts to generate complete *Orientia tsutsugamushi* genomes by whole genome sequencing have been limited by the difficulties of sequencing and assembling a repeat-dense genome, and no further genomes have been completed since the Boryong and Ikeda genomes in 2008. Current draft assemblies are fragmented with over 50 contigs per genome, and vary in size – the two assemblies of the genome of *Orientia tsutsugamushi* str. Karp available on Genbank are 1,459,958bp (https://www.ebi.ac.uk/ena/data/view/LANM01000000) and 2,022,909bp (https://www.ncbi.nlm.nih.gov/nuccore/LYMA00000000; Liao et al., 2017) in length, suggesting that assemblies are either incomplete, or have problems caused by the misassembly of repeats or the inclusion of contaminating sequences.

In this work, we have used Pacific Biosciences long-read sequencing to assemble six complete genomes of *Orientia tsutsgamushi* strains representing a range of geographical origins and serotypes. From this, we gain new insights into potential mechanisms underlying the characteristic antigenic diversity of Orientia, which may contribute to its widespread prevalence among humans. Finally, this expanded genomic perspective will contribute to our understanding of the phylogeography and epidemiology of this species, as well as contribute to more detailed studies of virulence mechanisms.

## Methods

### Bacterial propagation

All experiments were performed using *O. tsutsugamushi* grown in the mouse fibroblast cell line L929. Uninfected L929 cells were grown in 25 cm^2^ and 75 cm^2^ plastic flasks at 37 °C and 5% CO_2_, using DMEM or RPMI 1640 (Thermo Fisher Scientific) media supplemented with 10% FBS (Sigma) as described previously (Giengkam et al 2015). Infected L929 cells were grown in the same way, but at 35 °C. Frozen stocks of bacteria were grown for 5 days, then the bacterial content was calculated using qPCR against the bacterial gene TSA47 (Giengkam et al., 2015). Bacteria were isolated onto fresh L929 cells in 75 cm^2^ flasks at an Multiplicity of Infection of 10:1 and then grown for an additional 7 days. At this point bacteria were isolated from host cells and prepared for DNA extraction.

### DNA extraction

The supernatant was removed from infected flasks and replaced with 6-8 ml pre-warmed media. Infected cells were harvested by mechanical scraping and then lysed using a bullet blender (BBX24B, Bullet Blender Blue, Nextadvance USA) operated at power 8 for 1 min. Host cell debris was removed by centrifugation at 300xg for 3 minutes, and the supernatant was filtered through a 2.0 µm filter unit. 10 µl of 1.4 µg/µl DNase (Deoxyribonuclease I from bovine pancreas, Merck, UK) was added per 1 ml of bacterial solution, then incubated at room temperature for 30 minutes. This procedure removed excess host cell DNA. The bacterial sample was then isolated by centrifugation at 14,000xg for 10 min at 4 °C, and washed two times with 0.3M sucrose (Merck, UK). After the washing steps were completed DNA was extracted using a QIAGEN Dneasy Blood & Tissue Kit (QIAGEN, UK) following the manufacturer’s instructions.

Purified DNA samples were analysed by gel electrophoresis using 0.8% agarose gel, in order to assess the DNA integrity. The yield of genomic DNA was quantified using a nanodrop (Nanodrop^TM^ 2000, Thermo Scientific, UK) and Qubit Fluorometric Quantitation (Qubit^TM^ 3.0 Fluorometer, Thermo Scientific, UK).

### Sequencing

SMRTBell templates were prepared from purified *Orientia* genomic DNA according to PacBio’s recommended protocols. Briefly, 20kb libraries were targeted; enrichment for large fragments was done using BluePippin (Sage Science) size selection method or successive Ampure (Beckman Coulter) clean-ups, depending on the original DNA size distribution and quantity, as recommended by PacBio. SMRTBell templates were sequenced on a Pacific Biosciences RSII Sequencer using P6 chemistry with a 240min run time. An average of 1.05 Gb of raw sequence was collected per strain (range 0.3-2.4 Gb), with an average N50 read length of 28.5 Kb (range 10.6-41.5 Kb). Genomes were assembled using the RS_HGAP_Assembly.3 protocol from the PacBio SMRTPortal (version 2.3.0), with initial polishing performed on trimmed initial assemblies using the same raw sequencing data with the RS_Resequencing.1 protocol. Each assembly was further polished using paired-end reads sequenced on an Illumina Miseq machine. Sequencing information and Illumina data availability for each sample can be found in Table S1; PacBio data is available under EBI accession PRJEB24834. For each assembly, the corresponding Illumina reads were aligned to the PacBio assembly using Stampy v1.0.23 (Lunter and Goodson, 2010). Pilon v1.16 (Walker et al., 2014) was then used to generate a final genome, and corrected 2 to 265 errors in the assemblies, with the majority of the errors being single base deletions at the end of A or T homopolymer runs. All genomes were rotated and reverse complemented as needed so that the predicted start codon for the dnaA gene formed the first nucleotide in the genome sequence. Sequencing and assembly statistics can be found in Table 2.

**Table 1.** Bacterial strains used in this study.

| Strain | Original Source | Source in this study | Reference |
| --- | --- | --- | --- |
| Karp | New Guinea, human patient, 1943 | Naval Medical Research Centre (NMRC) | (Enatsu et al., 1999) |
| Kato | Niigata, Japan, human patient, 1955 | NMRC | (Enatsu et al., 1999) |
| Gilliam | Indian-Burmese border, human patient, 1943 | NMRC | (Rights and Smadel, 1948) |
| TA686 | Thailand, animal (*Tupaia glis*), 1963 | NMRC | (Enatsu et al., 1999) |
| TA763 | Thailand, animal (*Rattus rajah*), 1963 | NMRC | (Enatsu et al., 1999) |
| FPW1038 | Thailand-Burmese border, human patient (pregnant), 2010 | Mahidol-Oxford Research Centre (MORU) | (McGready et al., 2010) |
| UT76 | Udon Thani, Thailand, human patient, 2003 | MORU | (Blacksell et al., 2008) |
| UT176 | Udon Thani, Thailand, human patient, 2004 | MORU | (Paris et al., 2009) |

**Table 2.** Assembly statistics for the 10 assemblies used in this analysis. Genomes marked with * are previously-assembled reference strains.

| Strain | Genome length<br>(bp) | Contigs | GC<br>percentage | Errors corrected by Illumina<br>sequencing |
| --- | --- | --- | --- | --- |
| Boryong* | 2,127,051 | 1 | 31 | - |
| Ikeda* | 2,008,987 | 1 | 31 | - |
| FPW1038 | 2,035,338 | 25 | 31 | 265 |
| Gilliam | 2,465,012 | 1 | 31 | 7 |
| Karp | 2,469,803 | 1 | 31 | 48 |
| Kato | 2,319,449 | 1 | 31 | 5 |
| TA686 | 2,254,485 | 1 | 31 | 28 |
| TA763 | 2,089,396 | 8 | 31 | 88 |
| UT76 | 2,078,193 | 1 | 30 | 2 |
| UT176 | 1,932,116 | 1 | 30 | 13 |

**Table 3.** Number of genes predicted in each strain after annotation with Prokka, and the number of genes which were annotated as hypothetical. The Boryong and Ikeda strains were reannotated with Prokka for consistency between strains.

| Strain | Genes | Annotated as hypothetical |
| --- | --- | --- |
| Boryong | 2443 | 547 |
| Ikeda | 2186 | 417 |
| FPW1038 | 2198 | 369 |
| Gilliam | 2709 | 463 |
| Karp | 2578 | 470 |
| Kato | 2406 | 465 |
| TA686 | 2546 | 474 |
| TA763 | 2212 | 396 |
| UT76 | 2247 | 420 |
| UT176 | 2086 | 325 |

The Boryong, Ikeda, and non-*Orientia* Rickettsial genomes used in this study were obtained from NCBI (Table S2).

The finished assemblies were annotated using Prokka v1.11 (Seemann, 2014), using a custom database created from the Boryong and Ikeda strains, which were previously annotated using the NCBI prokaryotic annotation pathway. The Boryong and Ikeda strains were re-annotated using Prokka for consistency with the other samples. Short gene names for all non-hypothetical gene products were checked manually (607 products). Where genes names were present for Boryong and/or Ikeda a consensus name based on these was selected. Where no short name was available, the long gene name was searched for in *E. coli* using the UniProt database, and where a single and unambiguous match was selected this was used. In cases of ambiguity the protein sequence from *Orientia* was used in a BLAST search against *E. coli, R. rickettsii* and *H. sapiens* and the short name of the closest match was selected. The key *Orientia* genes *TSA56, TSA47, TSA22, ScaA, ScaC, ScaD,* and *ScaE* were also manually annotated by taking known protein sequences from the UT76 strain and using BLAST to find the homologous genes in the other strains and give them the correct names. The single contig genomes were rotated to begin with the *DnaA* gene.

Repetitive regions of the genome were defined as regions of at least 1000bp in length which had a match with another 1000bp region with up to 100 differences (mismatches, insertions, and deletions) allowed. The repetitive regions were identified with Vmatch (Abouelhoda et al., 2004).

The core and accessory genome was identified using Roary (Page et al., 2015) with a threshold of 80% sequence identity required to consider two sequences part of the same gene group. Core genes were defined as genes present in every sample and as a single copy in every sample. The accessory genes identified using Roary were re-clustered using CD-Hit (Fu et al., 2012; Li and Godzik, 2006) with a cutoff of 80% identity across 95% of the length of the shortest protein to identify accessory genes which were truncated copies of other proteins. The correlation between gene order in each pair of samples was calculated by taking the order of the genes relative to the Karp strain and calculating the Spearman’s rank coefficient between each pair. COG categories were assigned using RPS-BLAST to find matches in the NCBI Conserved Domain Database (Marchler-Bauer et al., 2002) and assigning a COG category to these using cdd2cog (Leimbach, 2016). Core repeat genes were identified using protein clusters generated by CD-Hit to find gene groups which were present at more than 1 copy. The clusters were identified using CD-Hit on the proteins predicted by Prokka with a cutoff of 80% identity across 90% of the length of the shortest protein. Pseudogenes were identified from CD-Hit protein clusters where at least one protein was a truncated version of the longest protein in the group. As pseudogenes which are truncated at the 5’ end will not be annotated by Prokka, BLAST (Altschul et al., 1997) was using to screen for any additional pseudogenes in non-genic regions by searching for BLAST hits with protein identity >= 80% and an E-value <10^−15^. This method found a further 26-37 pseudogenes per strain.

Further analysis used BioPython (Cock et al., 2009) and the GenomeDiagram package (Pritchard et al., 2006). Figure 1 was created with Circos (Krzywinski et al., 2009). Statistical tests were carried out in R (R Core Team, 2014) and the Python SciPy library (Jones et al.). Phylogenies were inferred using Maximum Likelihood methods in RaxML (Stamatakis, 2014) under the GAMMA model of rate heterogeneity and bootstrap values calculated using the rapid bootstrap method. The input sequences were aligned with Clustal Omega (Sievers et al., 2011) (for the 56kDa/46kDa/MLST trees) or using the MAFFT alignments produced by Roary (for the core gene tree). Phylogenetic trees were drawn using the ape (Paradis et al., 2004) and phytools (Revell, 2012) R packages, and Robinson-Foulds distances were calculated using the phangorn (Schliep, 2011) R package.

**Figure 1:**
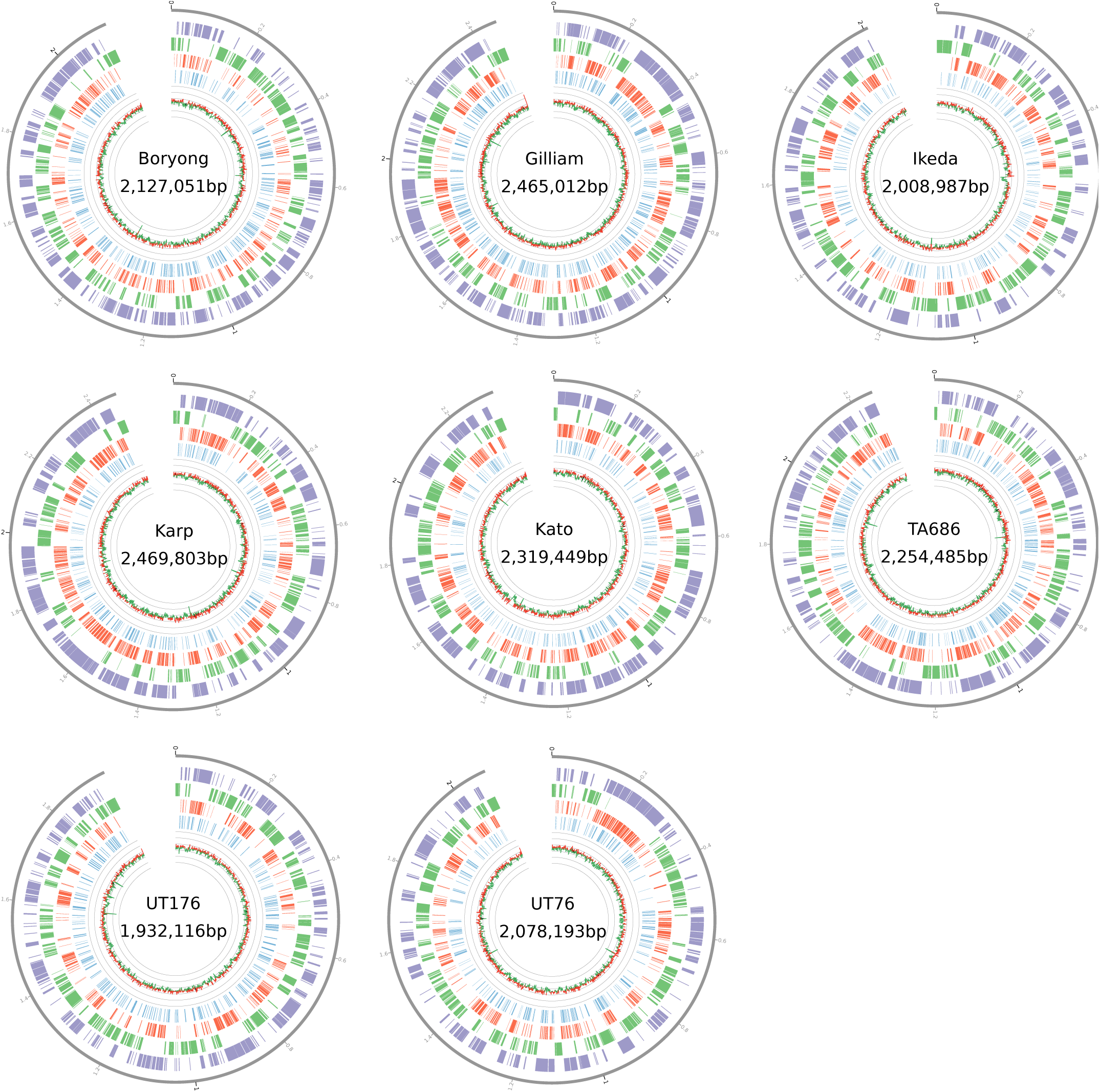
Ring diagrams for all single-contig strains. From outermost feature in each genome, moving inwards: repetitive regions are shown in purple, core genes in green, repeat genes in red and pseudogenes in blue. The track shows the GC percentage in windows of 1000bp. Values above the median GC are in green, and values below the median GC are in red.

**Figure 2:**
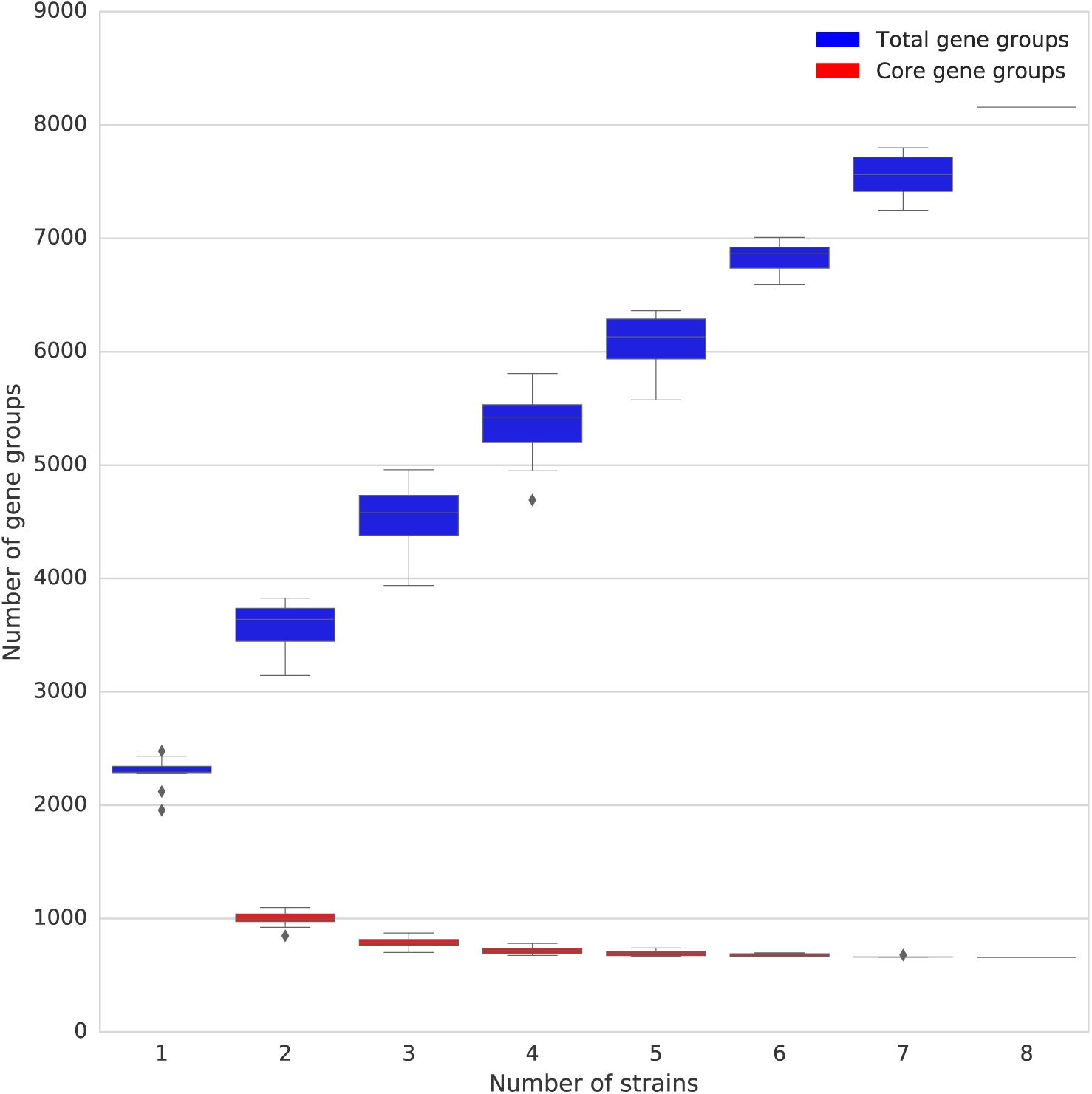
The number of core gene groups and the total number of gene groups (including the core gene groups) as more strains are added to the analysis. Boxplots represent all possible combinations of the number of strains given on the x-axis.

## Results

### Sequencing, Assembly, and Annotation

Eight genomes were assembled using the PacBio reads to perform initial genome assembly and Illumina sequencing reads to polish the genomes and reduce errors. Six of the eight genomes could be assembled into a single finished contig, while two genomes remain in multiple contigs. In addition, two previously assembled references genomes, *Orientia tsutsugamushi* str. Boryong and *Orientia tsutsugamushi* str. Ikeda, were incorporated into our analysis. The genome size ranges from 1.93Mb to 2.47Mb, and the GC content for all strains is consistent at 30-31%. We assessed the genomes to identify core genes shared between all genomes, and look for repetitive regions and repeat genes in each strain. Figure 1 plots the genetic elements of each complete genome.

The number of predicted genes in each strain ranges from 2086 (UT176) to 2709 (Gilliam) and is highly correlated with genome size (Spearman’s correlation coefficient 0.94, p < 2.2×10^−16^). A function could not be assigned, by similarity to reported sequences, to 325-547 genes (16 to 22 % of the identified coding regions) in each strain.

### Core genome analysis

The set of 8 complete, single-contig genomes was used to identify core genes (present in all genomes) and accessory genes (present in a subset of genomes), using the criterion that all members of a group of putative orthologues should be at least 80% identical (similar) to all other members of the group. While the unfinished genomes do not appear to have lower numbers of predicted genes, which might indicate the assembly is incomplete, for this analysis the two strains which assembled as multiple contigs were excluded to avoid excluding core genes which are missing from the unfinished assemblies. A total of 657 gene groups were present in all 8 strains and therefore form a putative core genome, while 2812 gene groups were present in 2-7 of the 8 strains, and a further 4687 gene groups were found in a single strain. The 657 core genes make up 28-35% of the genome of each strain (Table S3). The number of core gene groups does not continue to decrease as more genomes are added to the analysis, suggesting that the core genome of *Orientia* can be defined with 8 representative genomes. In the initial analysis with Roary, the total number of gene groups continues to grow, suggesting an open pan-genome, but observation of the 7499 accessory gene groups showed that of the 6050 groups where a function can be assigned to one or more gene, there are only 122 distinct gene products, many of them conjugal transfer proteins, transposases, DNA helicases, and other functions shared by genes known to be part of the *Orientia tsutsugamushi* amplified genetic element identified in the Ikeda strain (Nakayama et al., 2008). Re-clustering these accessory genes but allowing genes which are only a match to part of a gene sequence to cluster together to include more truncated and fragmented copies of genes shows that the number of accessory gene groups continues to increase, but at a slower rate (Figure S1). The number of gene products remains constant at 122 no matter how many strains are included in the analysis. This suggests that the increase in non-core gene clusters is mainly due to further duplication and truncation of existing genes, rather than by the import of novel genes.

### Genome Synteny

With the completed genomes produced by long read sequencing, the synteny of the genomes can be investigated. Previous work on the Boryong and Ikeda genomes showed extensive genome shuffling between the two strains. Analysis of the order and grouping of the core genes which are conserved in each genome shows that the genome has undergone massive rearrangement, with the core genes found in core gene ‘islands’ with repeat regions interspersed between these islands. The 657 core genes are present in 145-157 distinct islands, of which only 51 are conserved (defined as the same genes present in the same order) in all genomes. Figure 3 shows the position and ordering of these conserved core gene islands which are maintained in all samples relative to the position and ordering in the Karp strain. The correlation between gene order in each pair of samples is shown in Figure S2. A value close to 0 shows low correlation in gene order, while values closer to 1 show higher correlation in gene order. As there are differences in the correlation of gene order between strains, this suggests that the process of genome rearrangement is happening in multiple steps and not as a single event.

**Figure 3:**
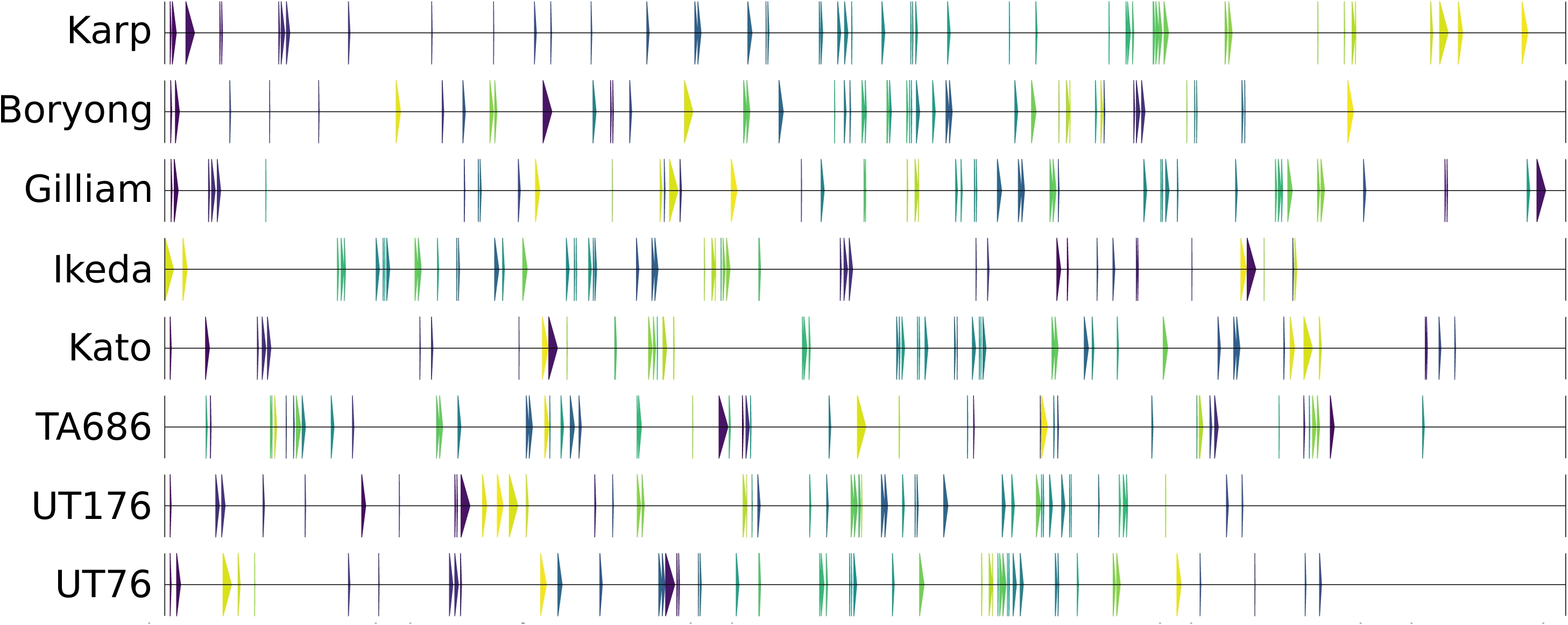
Each arrow represents the location of a core gene island containing one or more core genes which are conserved in the same order within an island across all strains. The arrows are coloured relative to their order in the Karp genome.

The identities of genes present on conserved islands is shown in Table S5. Conserved islands range from 1-13 genes in size, with larger islands often containing genes linked by plausible biological functions. For example, groups 3 and 6 include a number of cell division and peptidoglycan biosynthesis genes (including *mraY, murF, murE, pbp, ftsL, dnaJ* and *dnaK* in group 3 and *murC, murB, ddl* and *ftsQ* in group 6) and groups 31 and 32 include a number of 30S and 50S ribosomal proteins. Analysis of the number of conserved islands shared between samples shows that the number of conserved islands continues to decrease as more genomes are included (Figure S3), and suggests that gene order and clustering is not always constrained in *Orientia tsutsugamushi*. There is no difference seen in the size of the islands between conserved and non-conserved islands (Figure S4) (two-sample Kolmogorov-Smirnov test D=0.085, p-value=0.86), the nucleotide diversity between genes in the two categories of islands (Figure S5) (two-sample Kolmogorov-Smirnov test D=0.052, p-value=0.86), or the Clusters of Orthologous Groups (COG) categories assigned to genes in the two island categories (Chi-squared test χ^2^= 15.03, p=0.82).

### Repeats and pseudogenes

The genomes of *Orientia tsutsugamushi* are known to be highly repetitive, including a highly amplified genetic element known as the *Orientia tsutsugamushi* amplified genetic element (Otage), as well as other transposable elements.

Our results emphasise the large number of repeated genes and regions, including many genes related to the Type IV secretion system. The total proportion of the genome which is repetitive (see Methods for our definition of repetitive) differs markedly from 33% in UT176 to 51% in Gilliam (Table S3). In contrast, the extremely compact (and therefore non-repetitive) *Rickettsia typhi* genome is 0% repetitive by our measure and even, intriguingly, the *Rickettsia* endosymbiont of *Ixodes scapularis*, known to encode multiple copies of the same repetitive element found in *Orientia* (Gillespie et al., 2012), is 20% repetitive in our analysis, despite our methods giving more conservative figures than previously determined for the Ikeda strain (Nakayama et al., 2008).

We identified 530 groups of repeat genes containing 12043 genes present in multiple copies in at least one strain, which we term “core repeats”. Of the 530 groups, 427 represent genes found in multiple strains, which 103 are found only in a single strain. Despite clustering in 530 groups, the genes have only 66 different functional products, as is expected from the earlier results looking at all the non-core genes. The repeat genes are mainly transposase and conjugal transfer genes, similar to those previously reported in the Otage (Table S6), and cluster into genetic elements which are interspersed between the core genes. Many of these genes are present in high copy number, with all strains carrying over 200 transposases and 300 conjugal transfer genes and gene fragments. These core repeat elements occupy 35-47% of the *Orientia tsutsugamushi* genome and represent 57-67% of the genes in these genomes (Table S4).

*Orientia tsutsugamushi* genes are known to exhibit high levels of pseudogenisation and gene decay. We searched for pseudogenes in each genome, and identified up to 484 pseudogenes per strain (Table S7). This is lower than previously reported in Ikeda, but due to methodological differences the figures cannot be directly compared. We also assessed whether the pseudogene had been caused by truncation at the 5’ or 3’ end of the sequencing, or by frameshift.

### Phylogenetics

A phylogenetic tree was constructed using the core genes from each strain. This can be compared to trees built using the 56kDa (Figure 4) and 47kDa (Figure S6) genes, which are often used for phylogenetic analysis of *Orientia tsutsugamushi,* or to trees built using the MLST genes (Figure S7). *Orientia* strains are commonly based on their similarity to reference strains, either from phylogenetics or serology. Compared to the 56kDa tree, the core gene tree suggests the Kato and Ikeda strains are more closely related to the Karp, UT176, and UT76 strains than the TA686 and Gilliam strains (Figure 4). Robinson-Foulds distances between trees are shown in Table S8; for this small number of strains, the distance is lowest between the 47kDa tree and the core genome tree.

**Figure 4:**
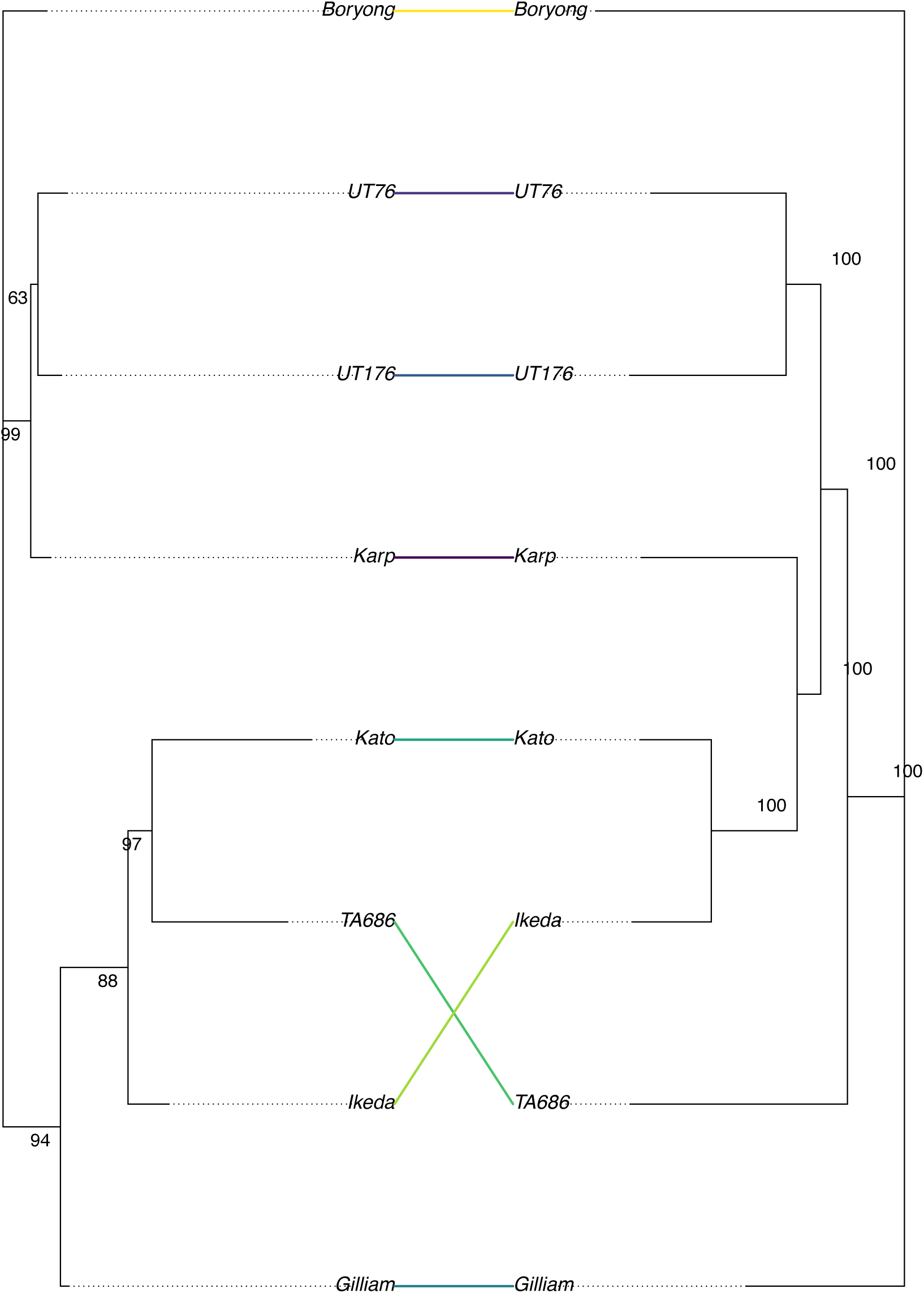
Phylogenetic trees generated from the 56kDa antigen sequence (left) and the sequence of the 657 core genes (right). The tree was inferred using the maximum likelihood method implemented in RaxML, and bootstrap values were calculated with the RaxML rapid bootstrap method.

## Discussion

We present the first large-scale study of *Orientia tsutsugamushi*, a bacterium which is important both for the study of human disease and for its unique insights into genome evolution.

Previous studies of *Orientia tsutsugamushi* genomes have used BAC cloning and Sanger sequencing to produce complete genomes (Cho et al., 2007; Nakayama et al., 2008), or have used next-generation sequencing strategies which have produced only incomplete and fragmented genomes (Liao et al., 2017). We demonstrate that a combination of PacBio and Illumina sequencing is sufficient to produce a single-contig genome, allowing us to study the gene content and synteny of this organism. For the two genomes which could not be assembled into single contigs in our study (FPW1038 and TA763), we found that the sequencing produced fewer reads at the high end of the length distribution. This suggests that given the highly repetitive nature of the *Orientia tsutsugamushi* genome, the DNA preparation and sequencing methods must be carefully chosen to produce very long reads in order to produce complete assemblies. We used Illumina sequencing to correct errors in our genomes, which was vital to reduce the number of homopolymer errors, which could otherwise suggest frameshift errors and affect gene annotation. While the fewest errors we corrected in a strain was two, this is likely an underestimate as errors in repetitive regions where Illumina reads cannot map are impossible to correct. While our analysis shows small differences when quantifying the extent of the repeat regions and repeat gene families in *Orientia* compared to previous work, a direct comparison is difficult due to differences in methodology between analyses.

Owing to the difficulties of producing complete genomes, most previous work has relied on single gene or MLST studies to investigate the genetic diversity of *Orientia tsutsugamushi.* We demonstrate that phylogenies generated from limited data are substantially different from those produced from the whole core genome. The common practice of grouping *Orientia* strains into ‘Karp-like’ or ‘Gilliam-like’ groups based on the genotype of the 56kDa antigen may not give an accurate view of the relatedness of these strains, especially when recombination is taken into account, although this may still be important when considering immune response.

Previous work has demonstrated limited synteny between the two reference strains of *Orientia tsutsugamushi*, but we extend this to demonstrate that there is minimal synteny between any known *Orientia tsutsugamushi* genome. The pattern of core gene islands separated by transposable elements and repeats suggests a repeat-mediated system of chromosome rearrangement. It is unclear whether this is a gradual process of genome rearrangement, or whether the genome is being broken apart and rearranged rapidly, similar to chromothripsis or the chromosome repair of *Deinoccocus radiodurans* after exposure to ionizing radiation. In *Deinococcus*, it is thought that RecFOR pathway is particularly important for DNA repair, and it has no homologues to RecB or RecC (Cox et al., 2010). Similarly, in *Orientia*, the core genome does not contain RecB or RecC, but does contain the RecFOR pathway genes, indicating this alternative DNA repair pathway may also be important. Longitudinal studies of *Orientia tsutsugamushi* genomes during passage or infection may be needed to determine the speed and processes of genome rearrangement in *Orientia*.

We report a core genome of only 657 genes, compared to the 519 previously reported as the core genome shared between *Orientia* and five other sequenced *Rickettsia*, despite using a relatively low sequence identity threshold to determine gene clusters. Differences in methodology may lead to the reporting of different core gene sets, but more interesting is the pattern of core genome islands separated by amplified repeat regions, and the lack of conservation in the ordering and clustering of the core genes.

All of the *Orientia* genomes show high repetitiveness, which we measured as both non-unique regions of the genome, and genes which are present in multiple copies (some of which may be truncated). The genomes of intracellular bacteria tend towards genome reduction and gene loss (Darby et al., 2007; Merhej and Raoult, 2011), but maintain degraded genes and accumulate non-coding DNA. The transition to intracellularity has been hypothesized to lead to the relaxation of selective pressure on the genome (Moran, 1996), with an increased rate of sequence evolution. The expansion of the Otage (and other mobile elements) throughout the *Orientia* lineage appears to be another consequence of relaxed selection on *Orientia* in its intracellular niche, again leading to accelerated sequence evolution of the genome through rearrangement and gene loss. This is supported by the finding that the diversity of gene repertoire between strains of *Orientia tsutsugamushi* is largely due to the duplication and truncation of existing genes, and we find no evidence for the acquisition of new genetic material via horizontal transfer. The amplication of a transposable element has been seen in Rickettsial (Gillespie et al., 2012) and non-Rickettsial (Wiens et al., 2008) species, but it is not known whether this is associated with rearrangement of the genome in other species.

In conclusion, we report the generation of six complete and a further two draft genomes from a diverse set of strains of the important but neglected human pathogen *Orientia tsutsugamushi.* This set includes the major reference strains Karp, Kato and Gilliam, and will serve as a valuable resource for scientists and clinicians studying this pathogen, in particular supporting future work on *Orientia* genomics, vaccine development, and cell biology. The new genomes reported here confirm the status of *Orientia* as one of the most fragmented and highly repeated bacterial genomes known, and exciting questions remain regarding the mechanisms and timeframes driving the evolution of these extraordinary genomes.

## Data Availability

Sequence data and assemblies generated in this study have been uploaded to the EBI under project PRJEB24834.

## Author Contributions

## Acknowledgements and Funding

J.S. was funded by a Royal Society Dorothy Hodgkin Research Fellowship. P.D. is supported by a Wellcome Trust Core Award (090532/Z/09/Z).

## Supplementary Figures and Tables

**Figure S1.**
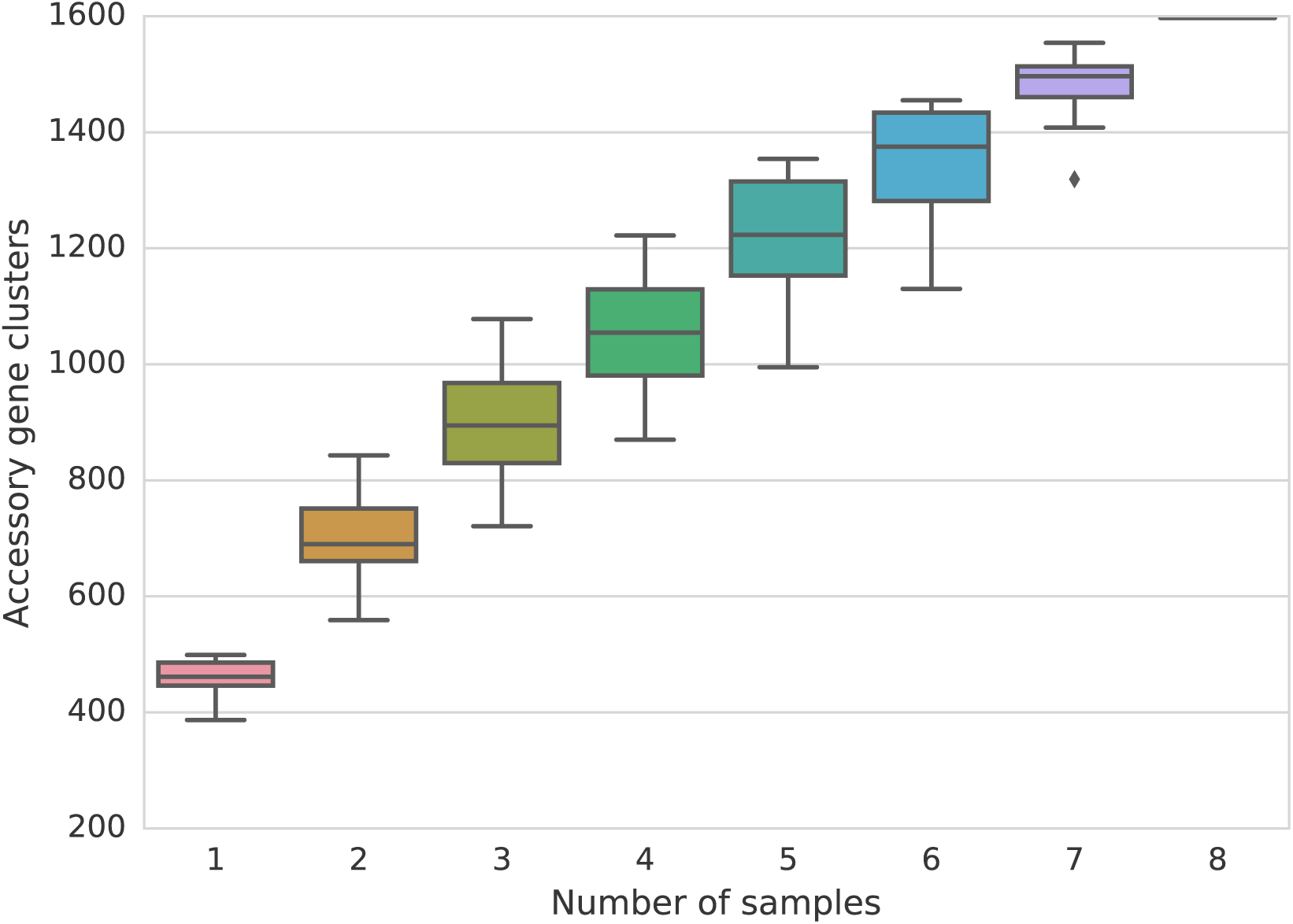
Boxplot of accessory genes clustered with a lenient length threshold to show how the number of clusters increases with number of samples included in the analysis.

**Figure S2.**
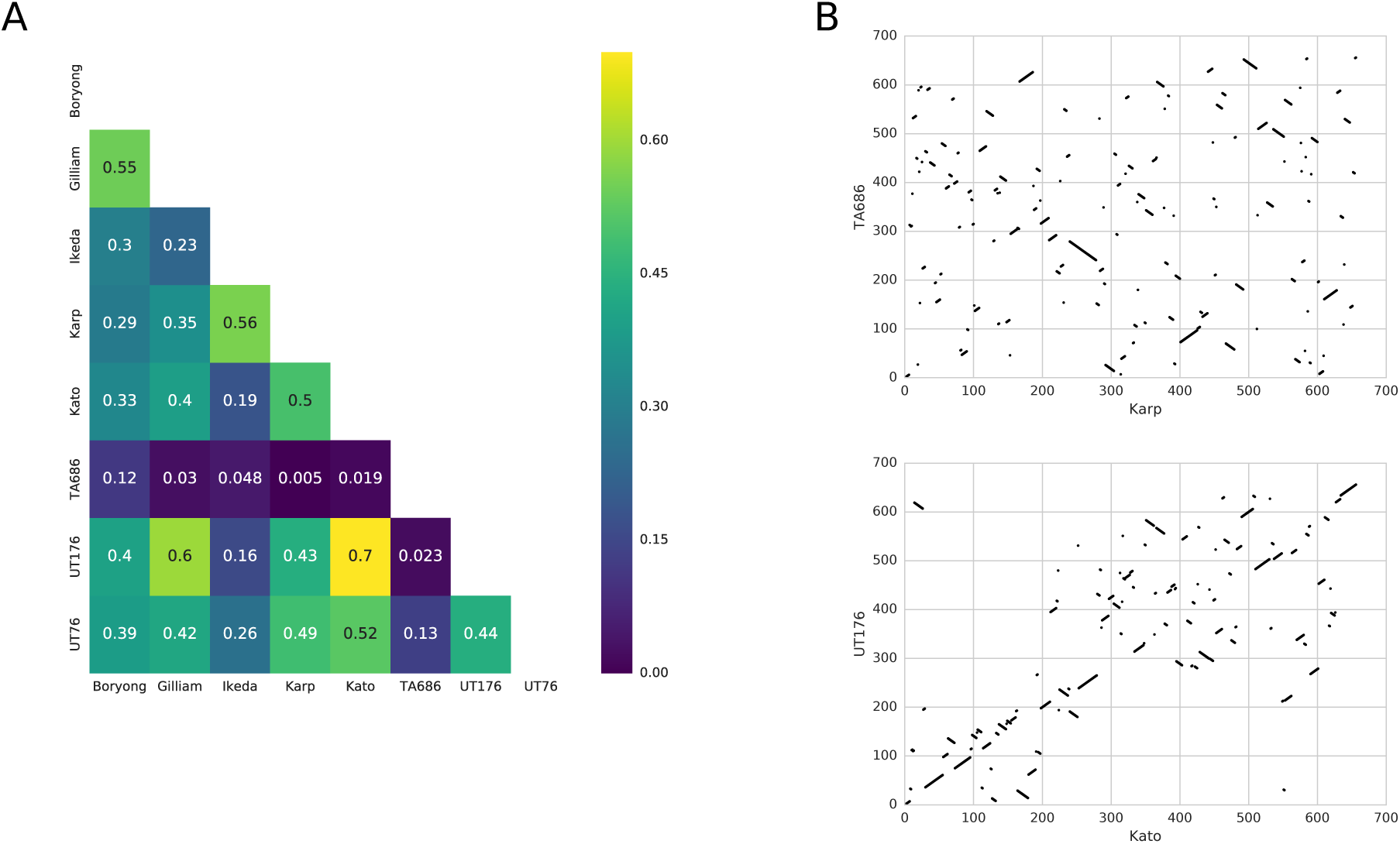
A - Heatmap showing the correlation in gene order between each pair of samples. B – dotplots showing the gene ordering between the pair with the highest correlation (Kato and UT176) and the lowest correlation (Karp and TA686).

**Figure S3.**
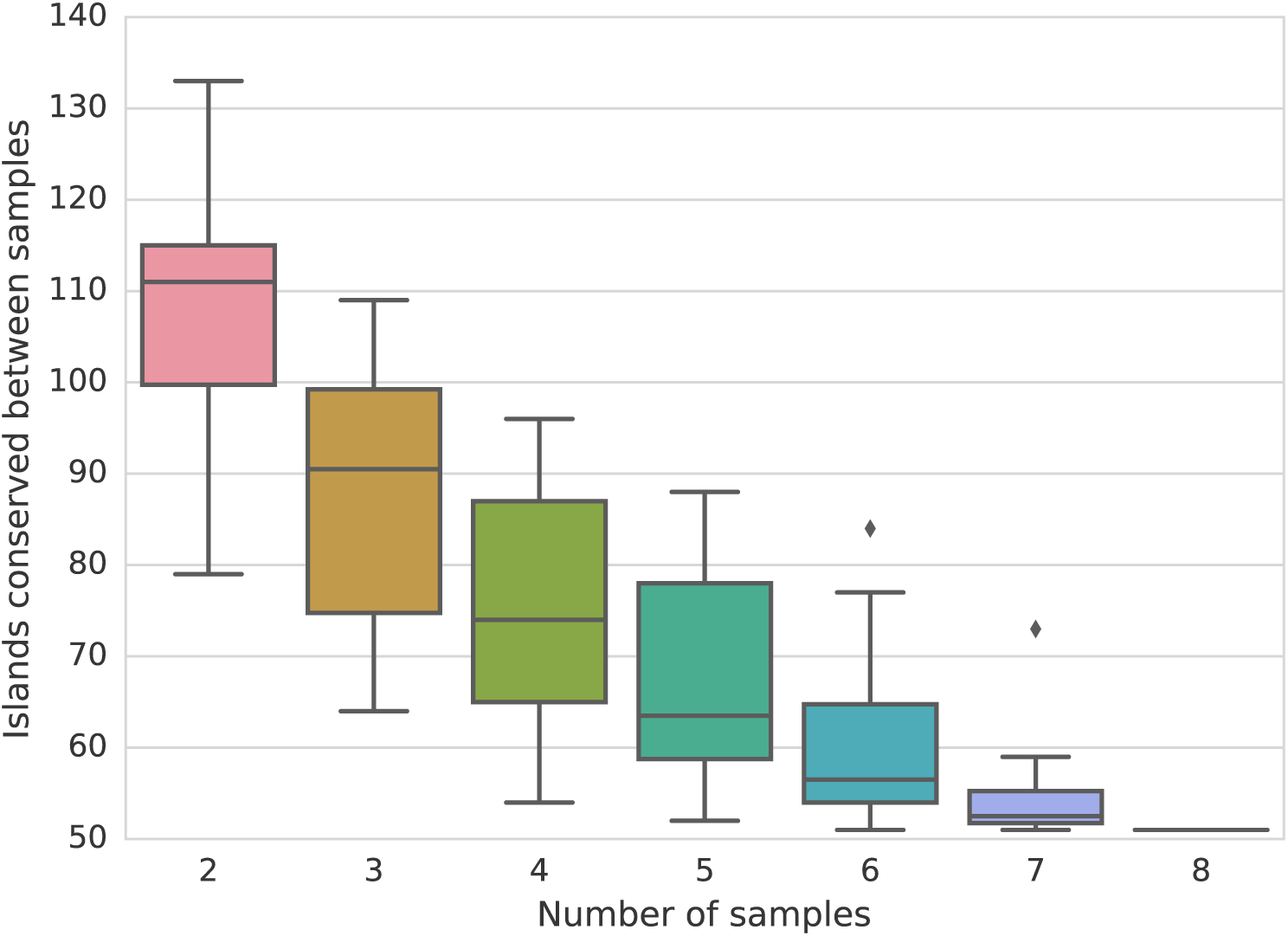
Boxplot showing the number of islands conserved between samples across all different combinations of samples.

**Figure S4.**
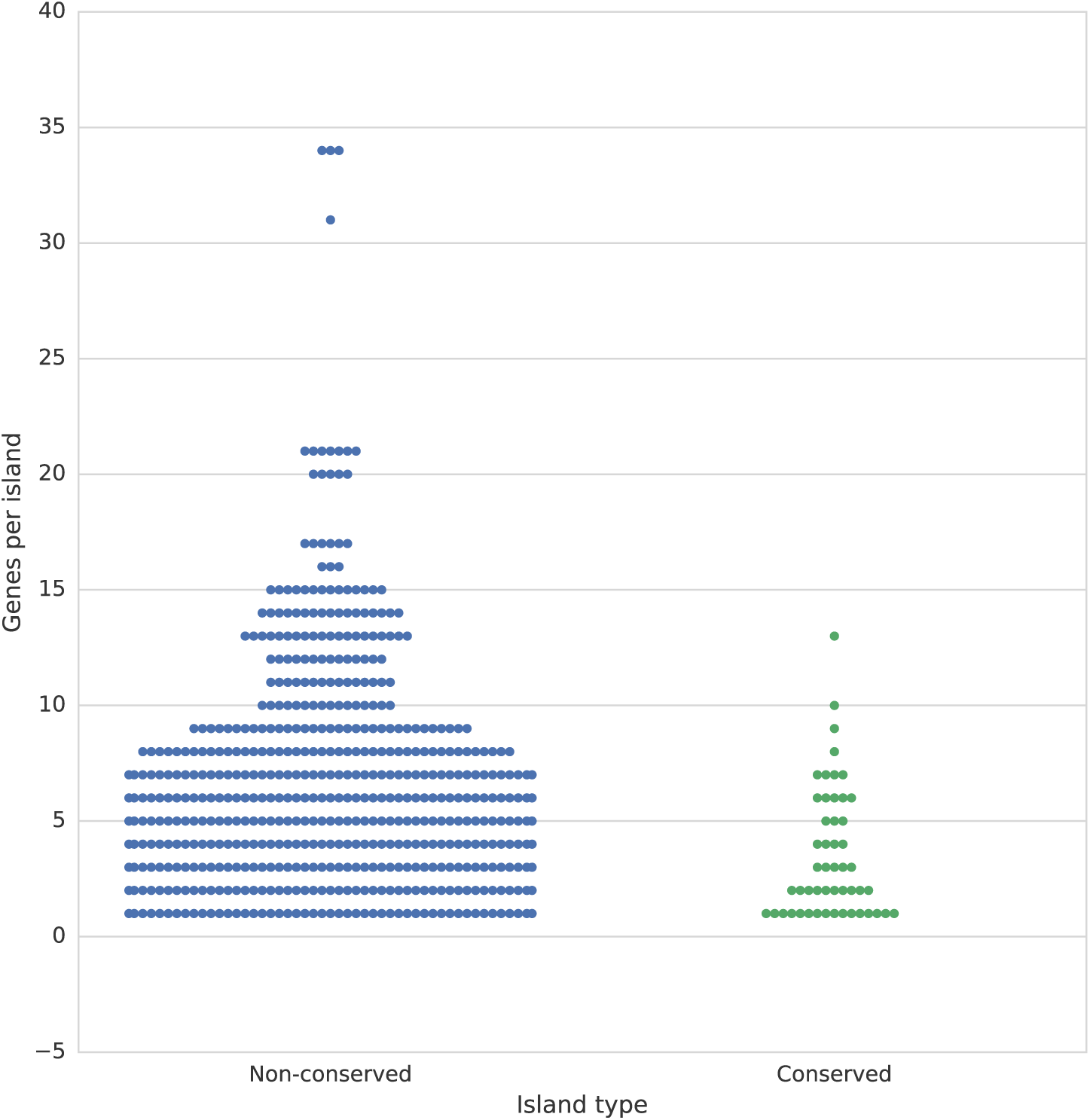
The number of genes per island in conserved versus non-conserved islands.

**Figure S5.**
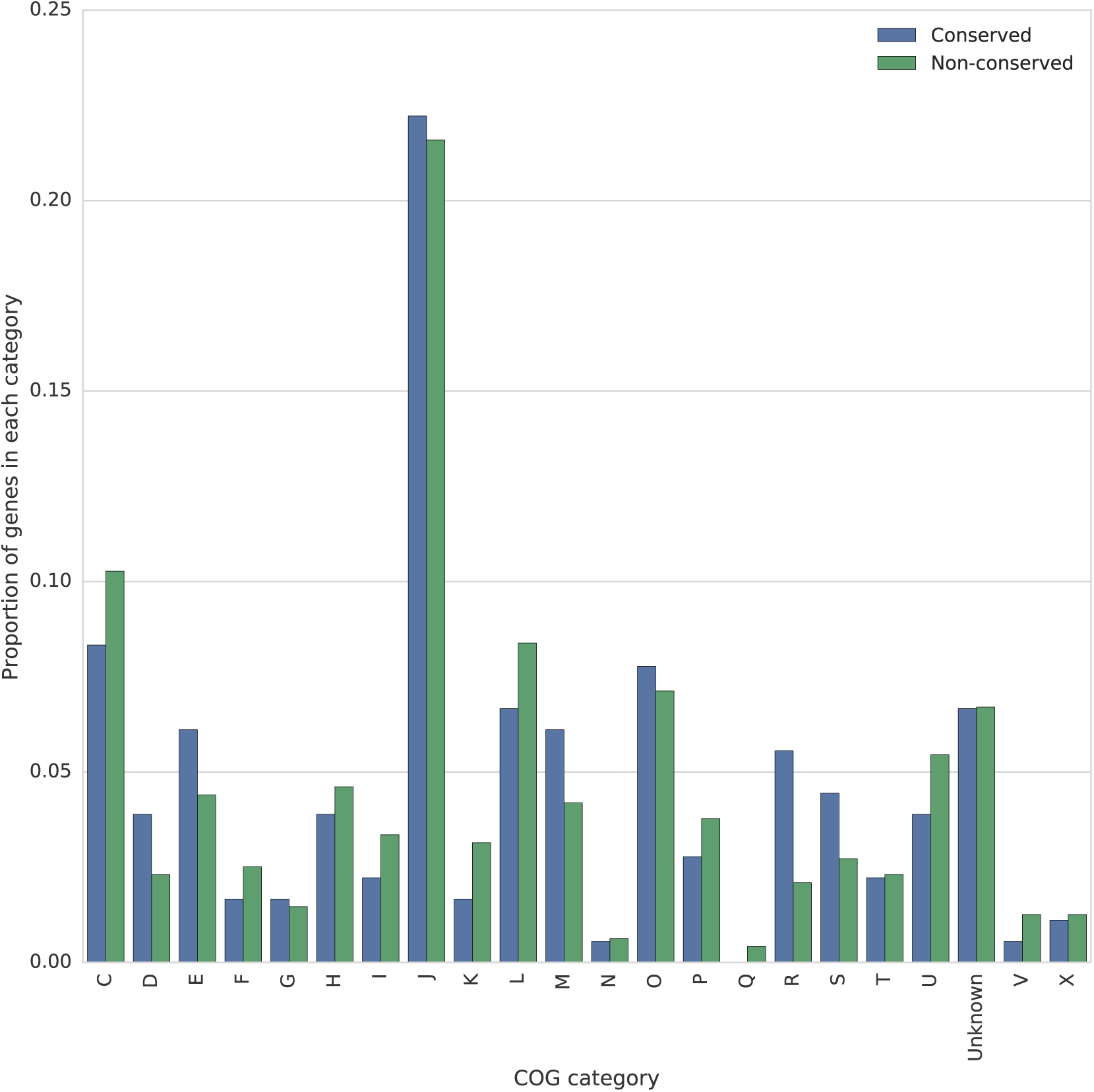
The proportion of core genes which are in conserved and non-conserved islands in each COG category.

**Figure S6.**
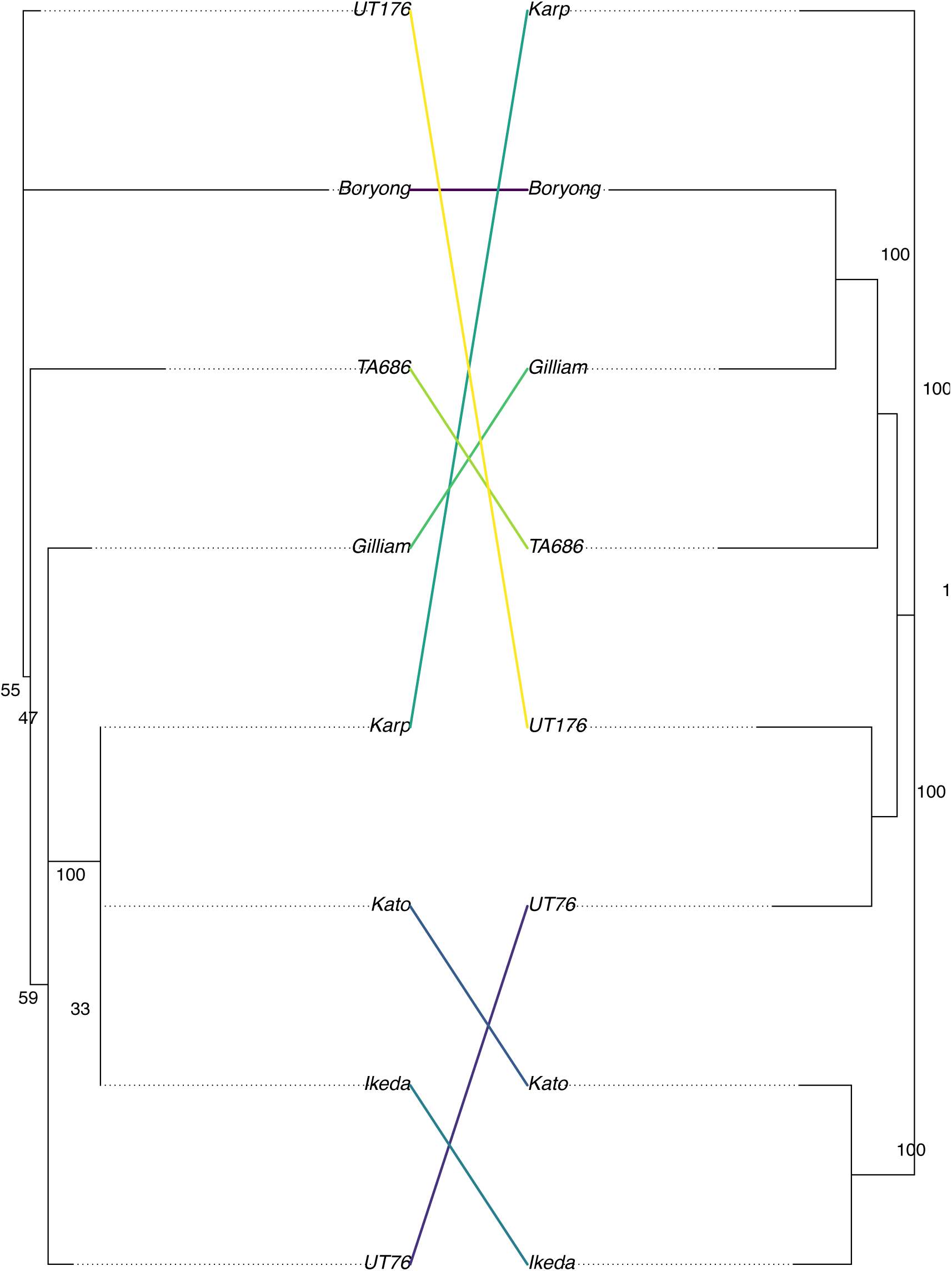
A phylogenetic tree showing the relationship between a tree generated using the 47kDa antigen sequences, and the sequences of 657 core genes.

**Figure S7.**
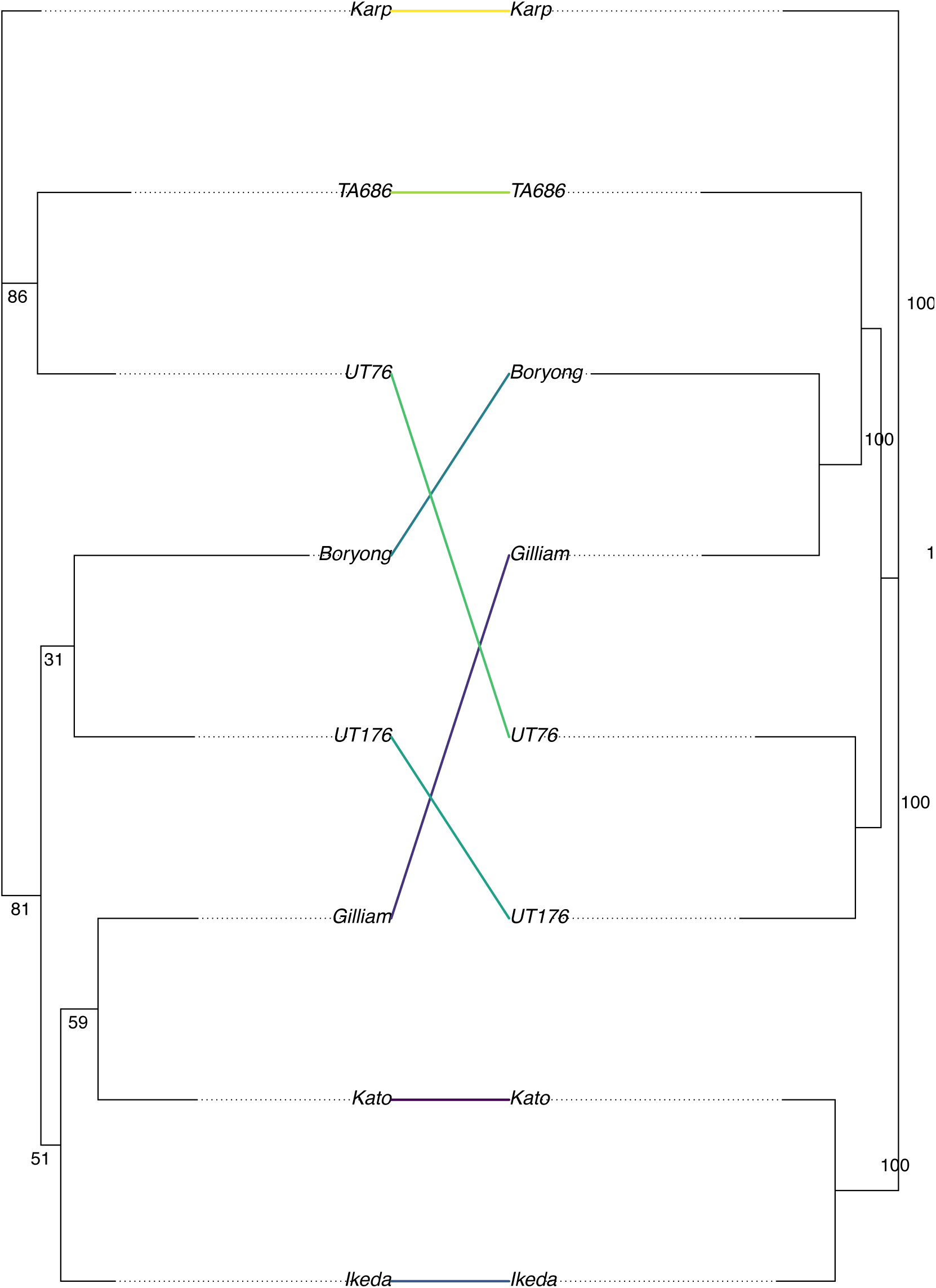
A phylogenetic tree showing the relationship between a tree generated using MLST gene sequences, and the sequences of 657 core genes.

**Table S1.**
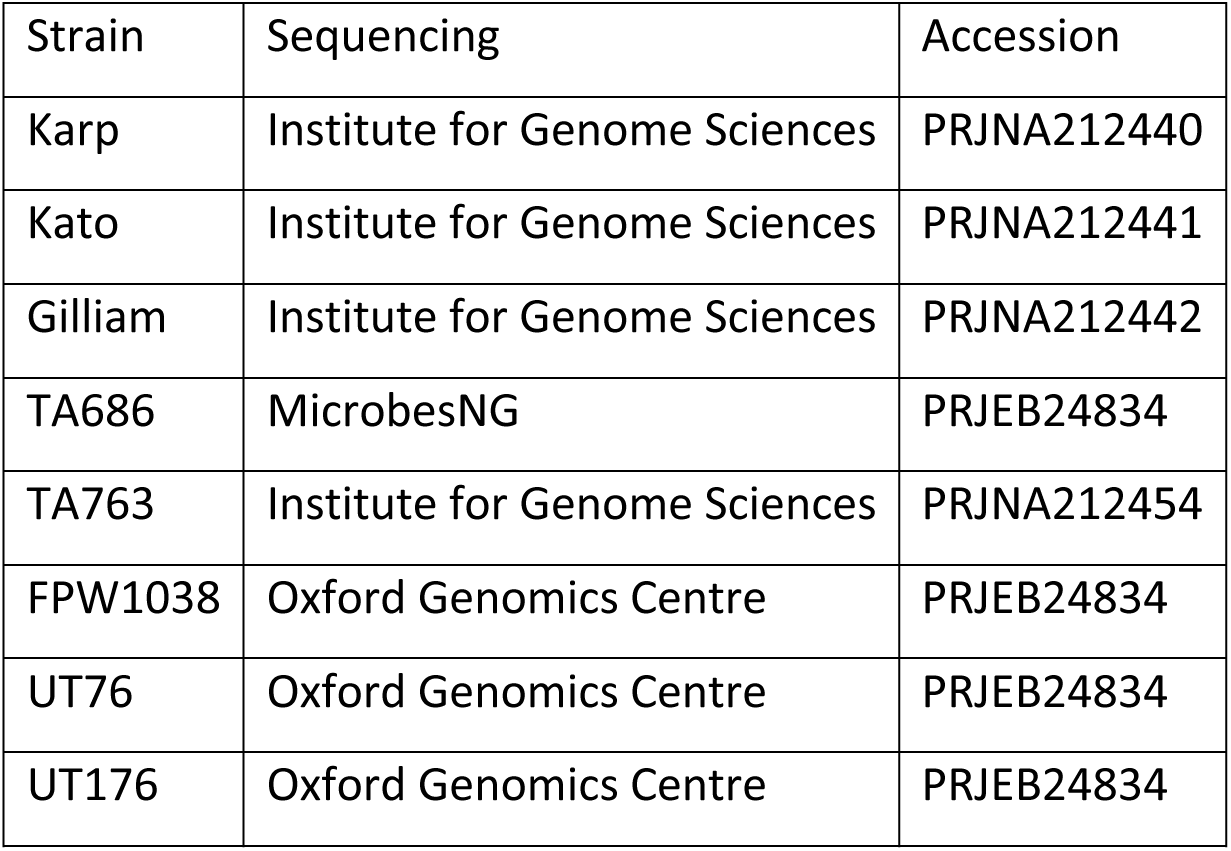
Sources and data accession for Illumina sequencing data.

| Strain | Sequencing | Accession |
| --- | --- | --- |
| Karp | Institute for Genome Sciences | PRJNA212440 |
| Kato | Institute for Genome Sciences | PRJNA212441 |
| Gilliam | Institute for Genome Sciences | PRJNA212442 |
| TA686 | MicrobesNG | PRJEB24834 |
| TA763 | Institute for Genome Sciences | PRJNA212454 |
| FPW1038 | Oxford Genomics Centre | PRJEB24834 |
| UT76 | Oxford Genomics Centre | PRJEB24834 |
| UT176 | Oxford Genomics Centre | PRJEB24834 |

**Table S2.** NCBI identifiers for previously published strains used in this paper.

| Genome | NCBI Identifier |
| --- | --- |
| *Orientia tsutsugamushi* strain Boryong | GCF_000063545.1 |
| *Orientia tsutsugamushi* strain Ikeda | GCF_000010205.1 |
| *Rickettsia typhi* strain Wilmington | GCF_000008045.1 |
| *Rickettsia* endosymbiont of *Ixodes scapularis* | GCF_000160735.1 |

**Table S3.**
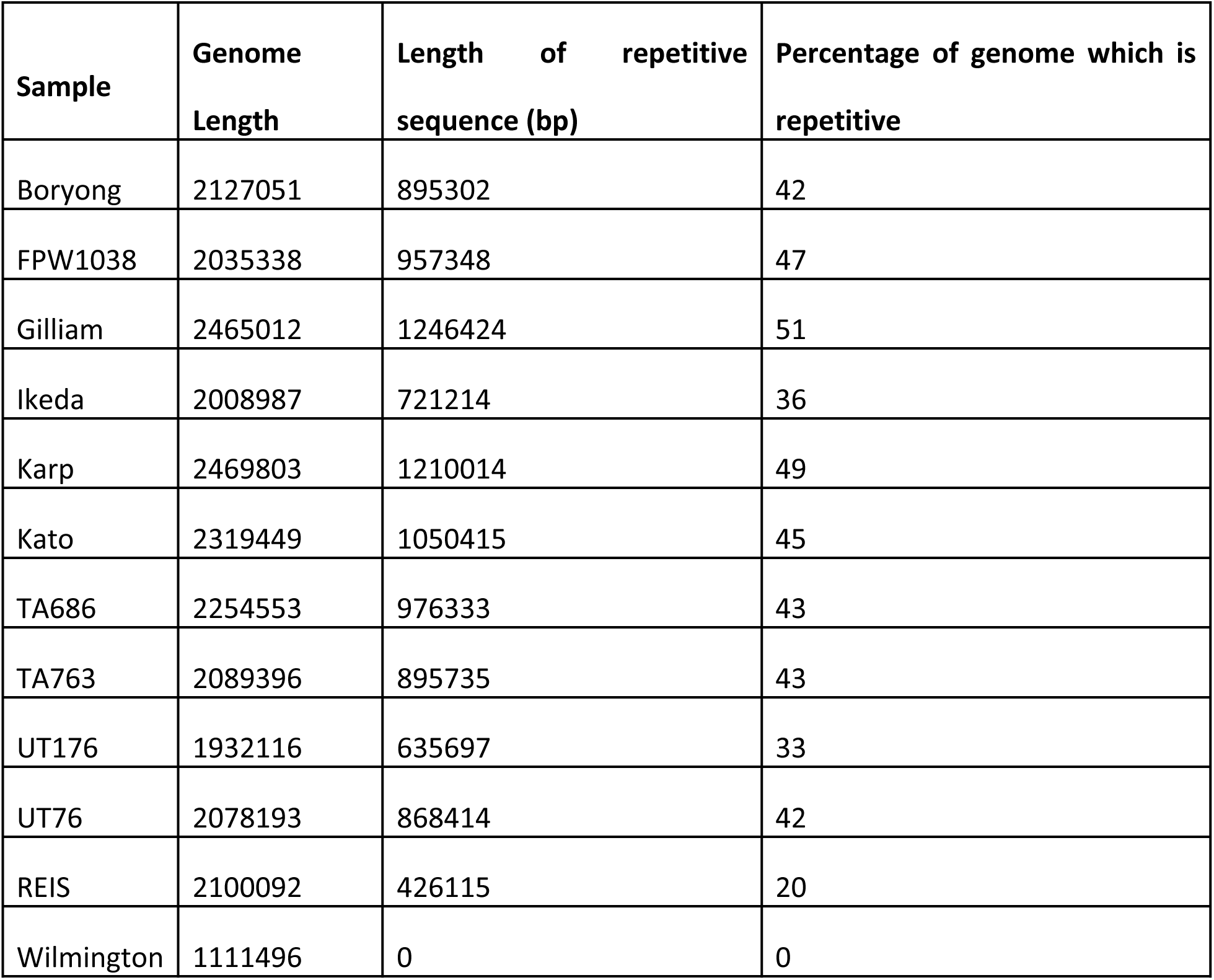
Total length of repetitive genome sequences in each strain, and as a percentage of the genome. REIS: Rickettsia endosymbiont of Ixodes scapularis. Wilmington: Rickettsia typhi strain Wilmington.

**Table S4.** Core gene and core repeat statistics.

| Sample | Genome Length | Length of core genes | Core genes as proportion of genome | Length of repeat genes | Repeat genes as percentage of genome |
| --- | --- | --- | --- | --- | --- |
| Boryong | 2127051 | 679631 | 0.32 | 748541 | 35 |
| Gilliam | 2465012 | 681491 | 0.28 | 1165831 | 47 |
| Ikeda | 2008987 | 683889 | 0.34 | 757868 | 38 |
| Karp | 2469803 | 682061 | 0.28 | 1163785 | 47 |
| Kato | 2319449 | 682142 | 0.29 | 1039243 | 45 |
| TA686 | 2254553 | 682706 | 0.30 | 933469 | 41 |
| UT176 | 1932116 | 681689 | 0.35 | 738572 | 38 |
| UT76 | 2078193 | 682964 | 0.33 | 826716 | 40 |

**Table S5.**
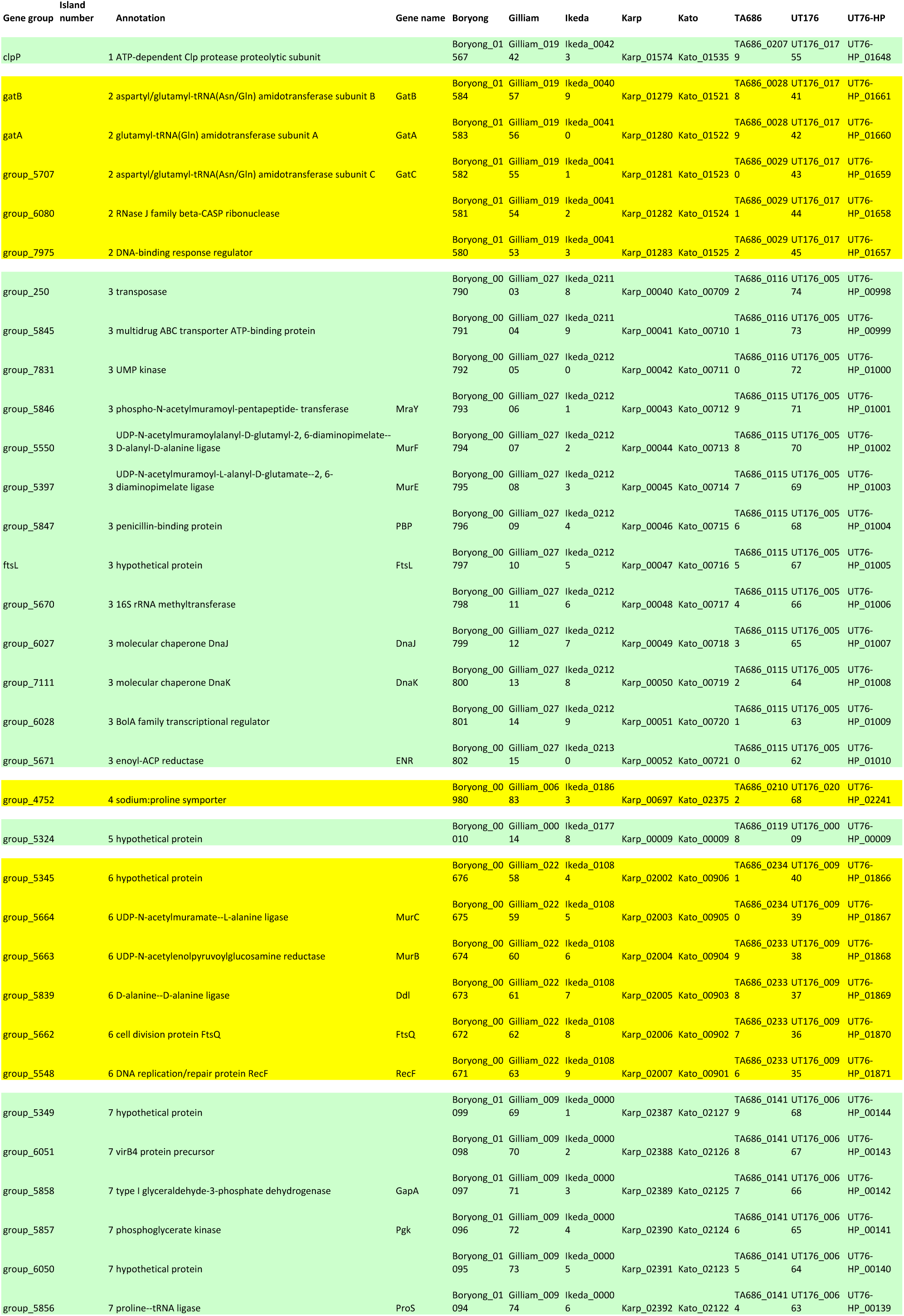
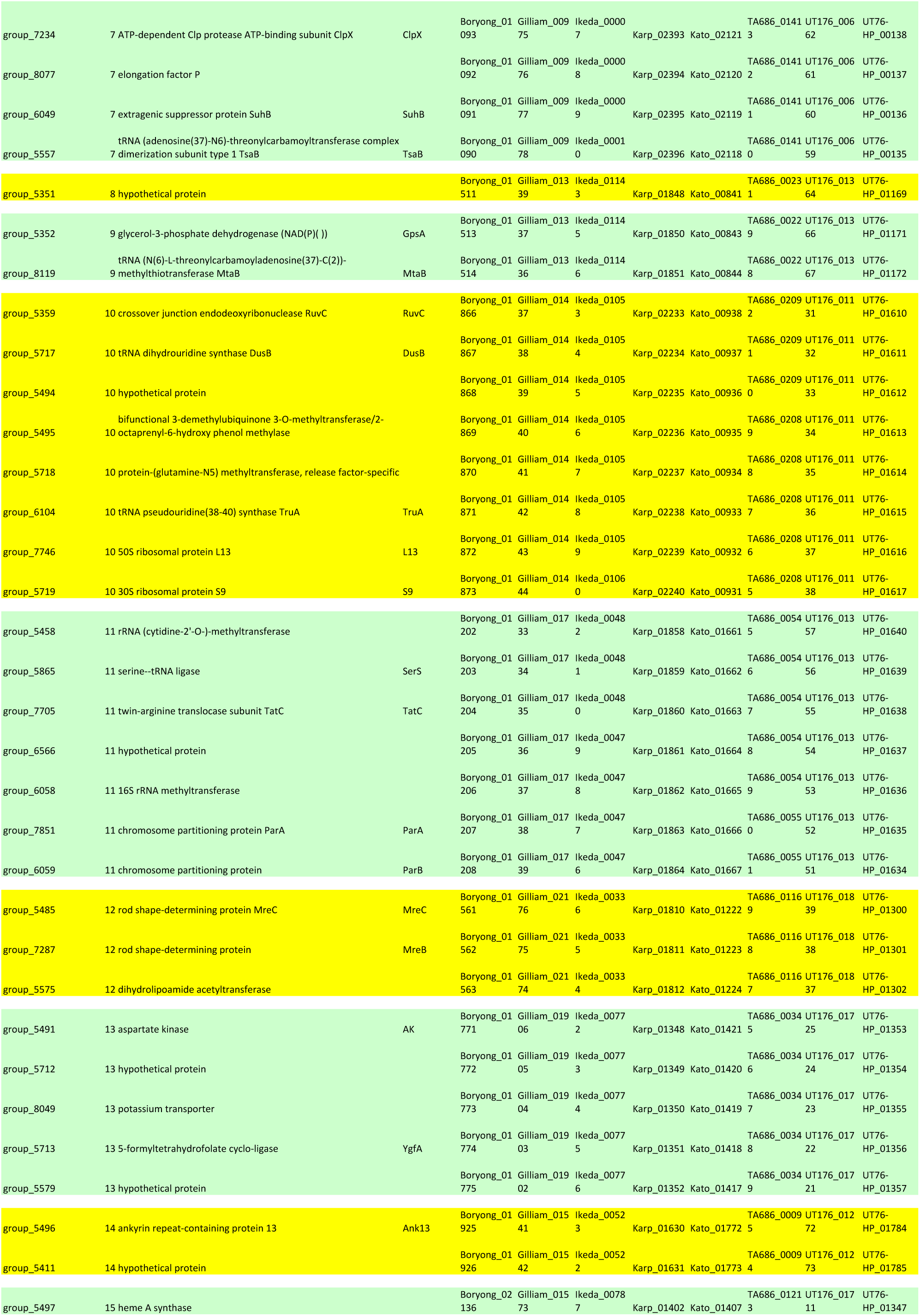
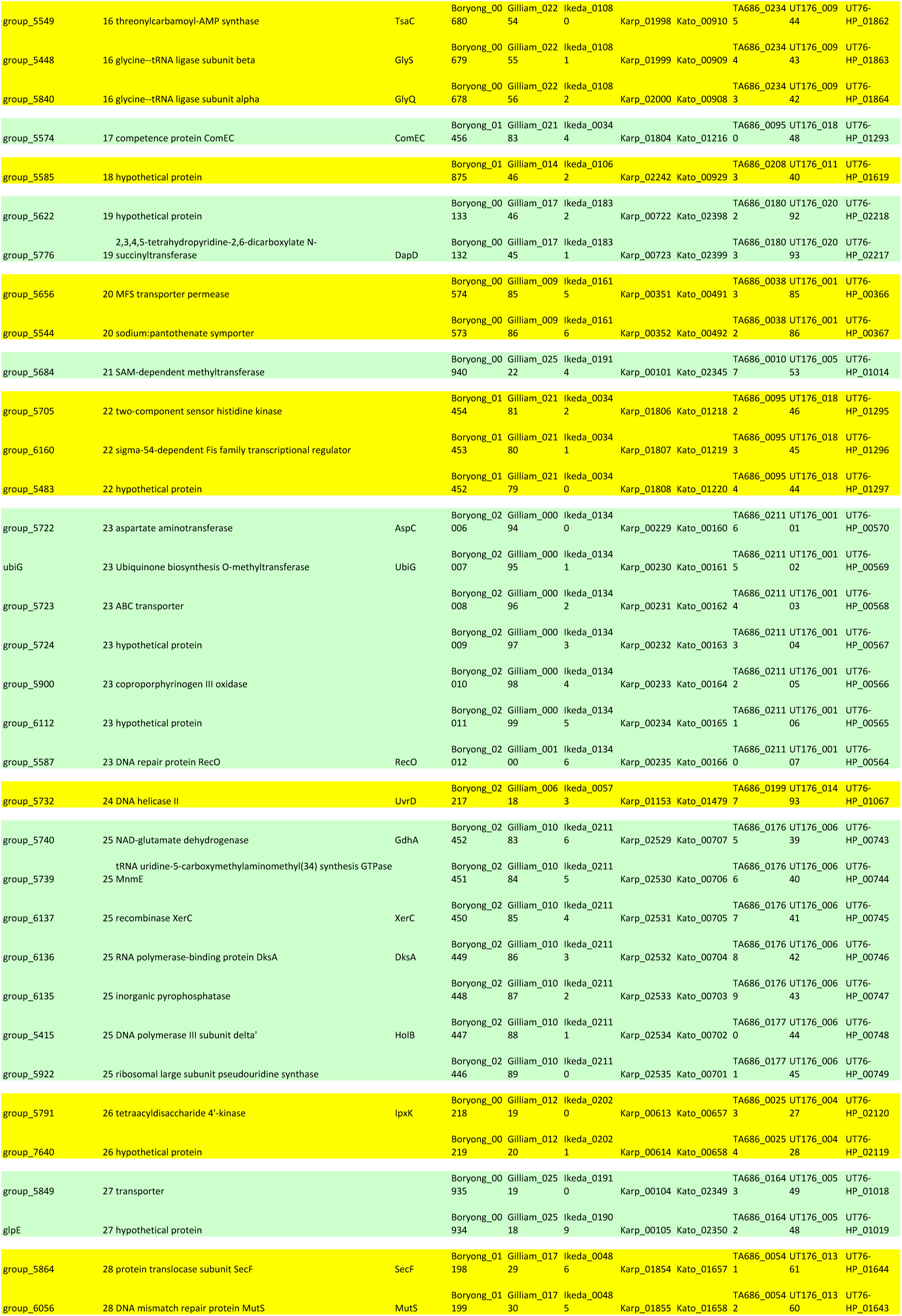
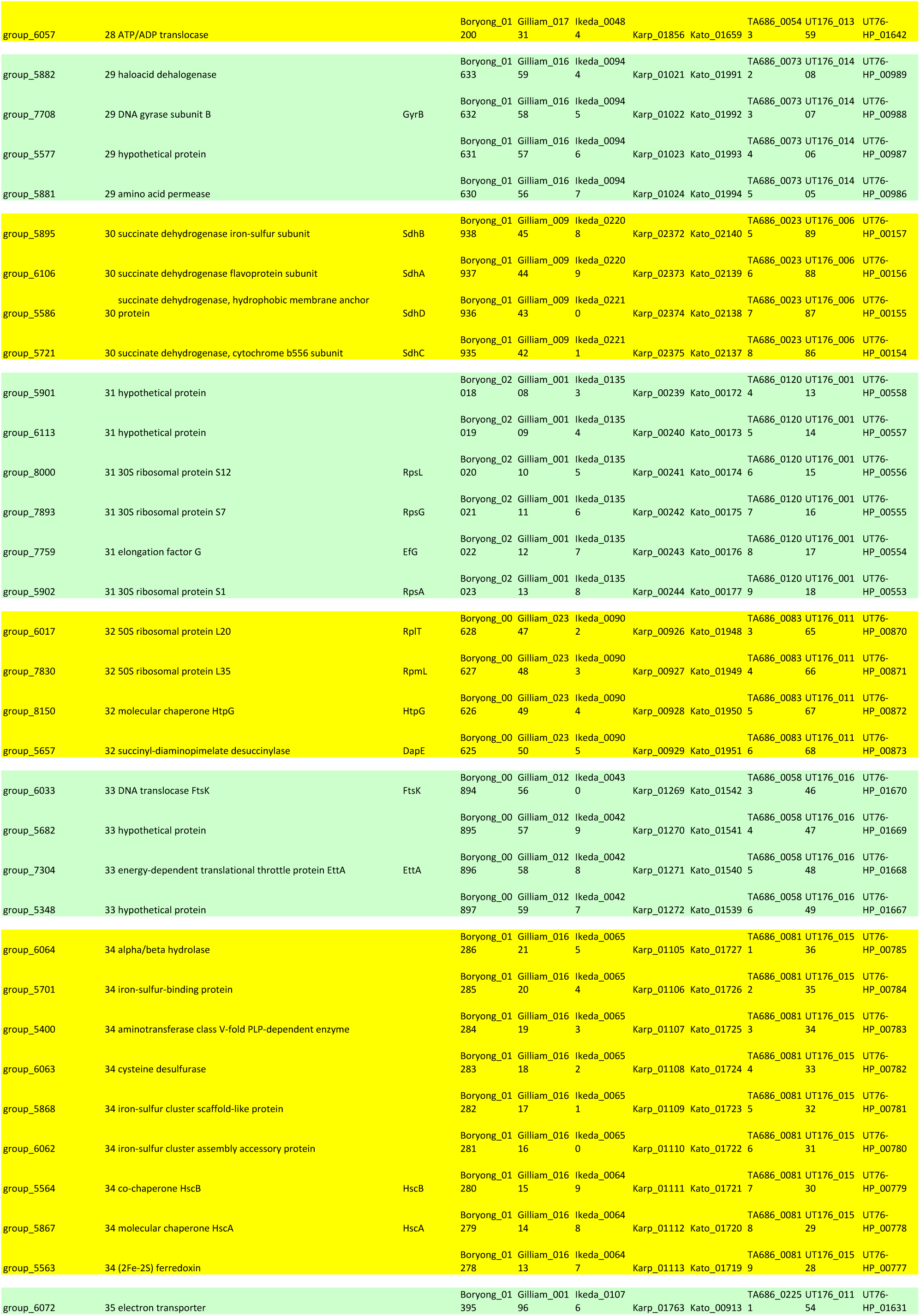
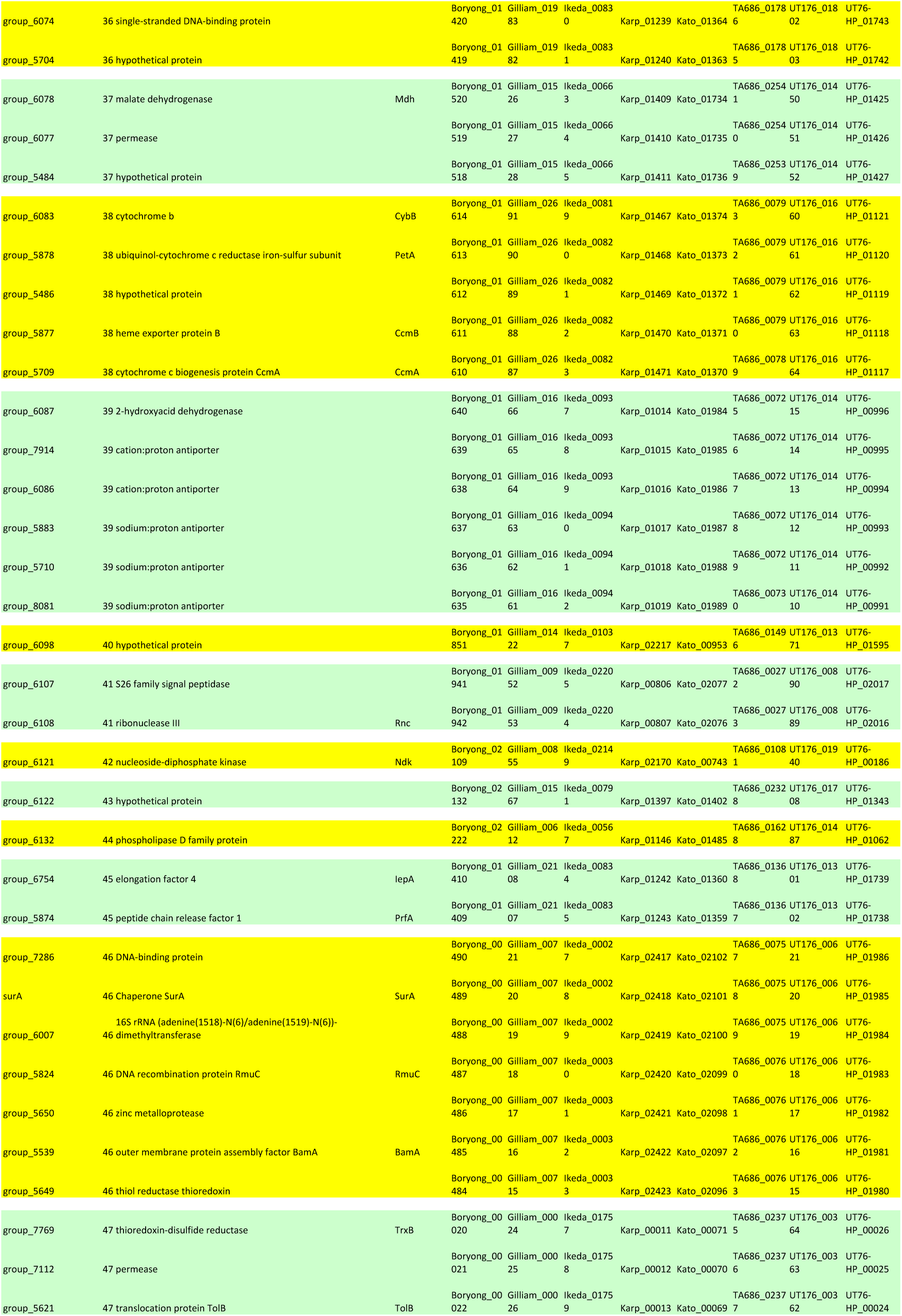
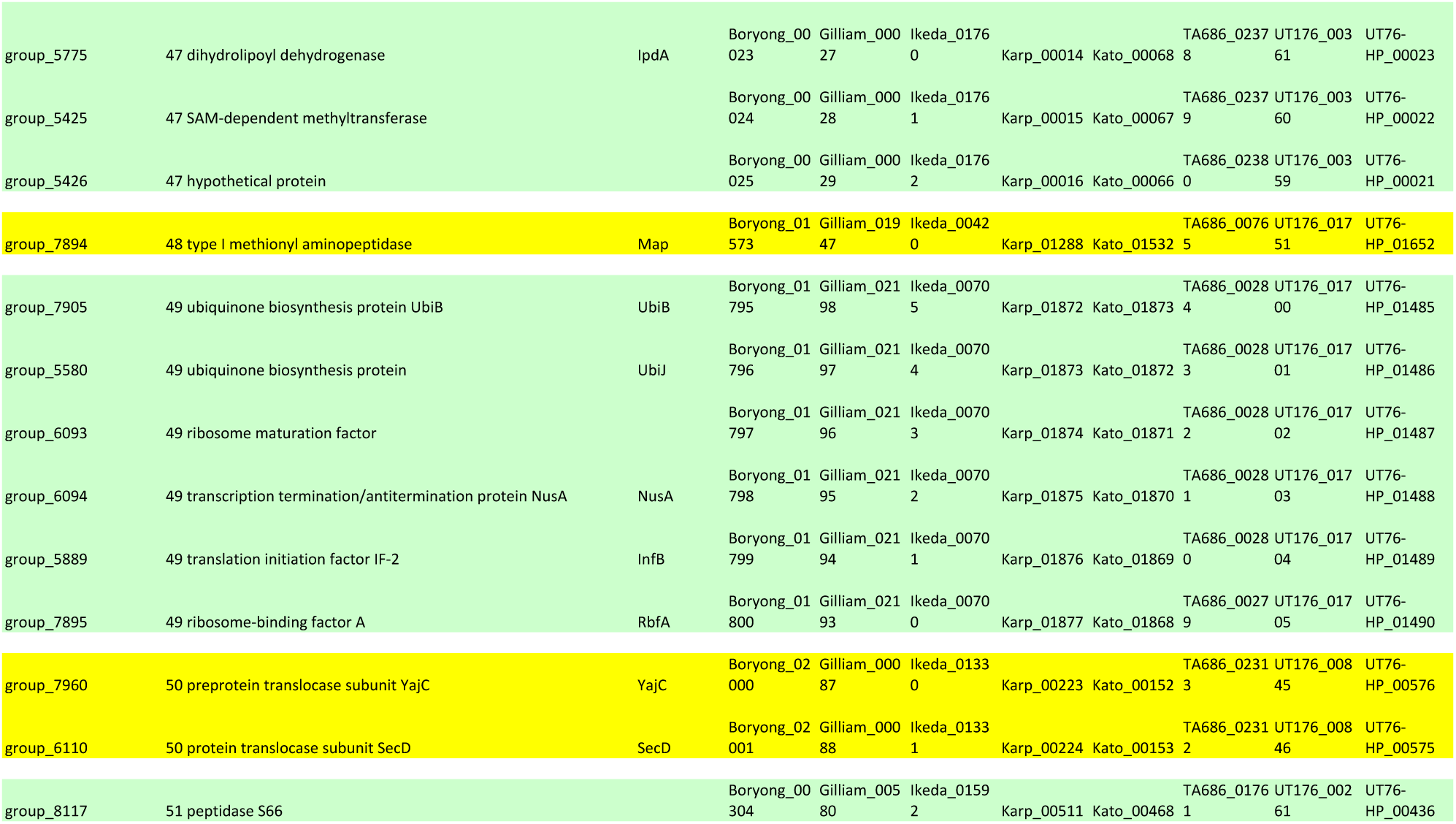
Core genes calculated by Roary. Gene names are given for the Karp strain.

**Table S6.**
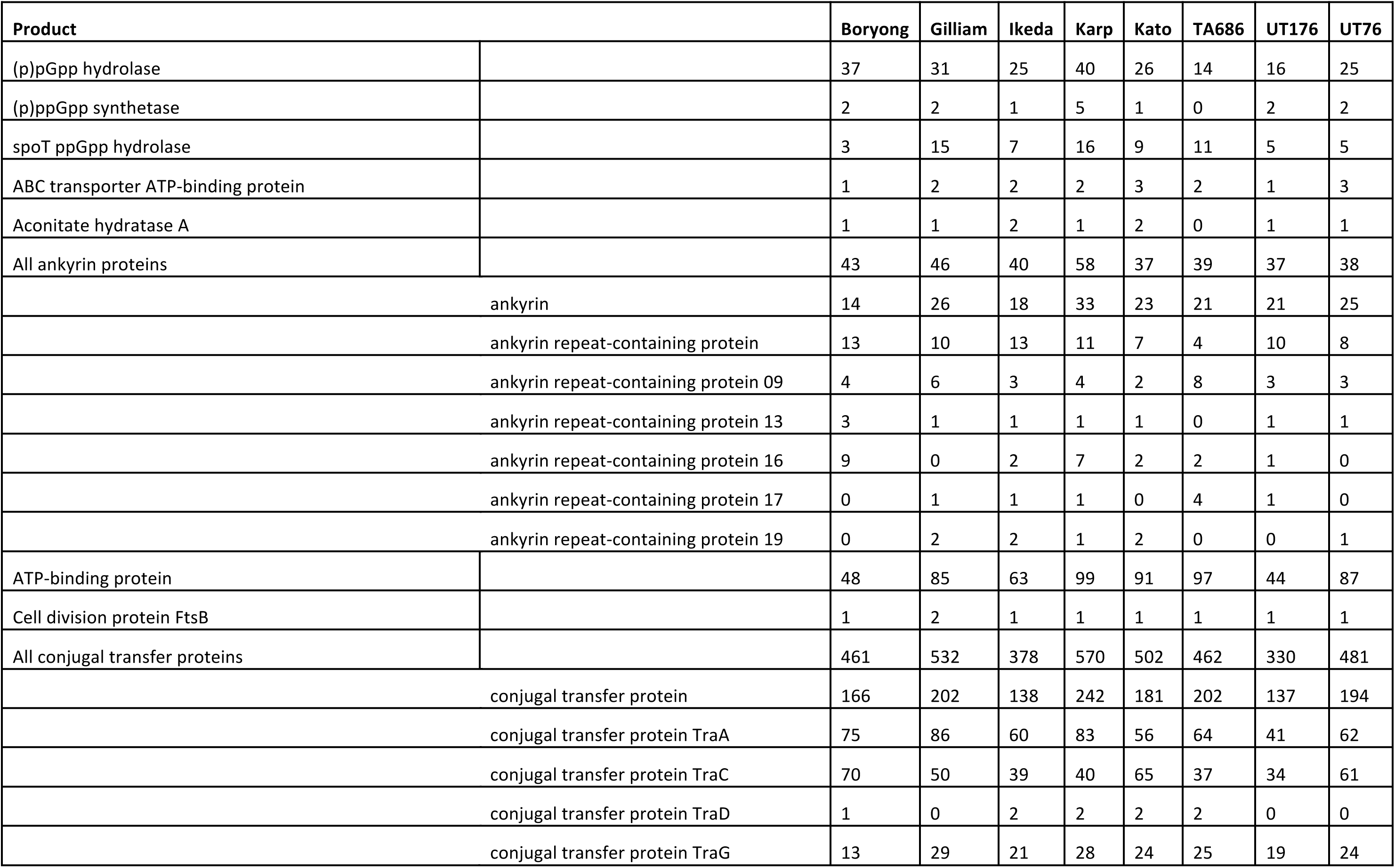
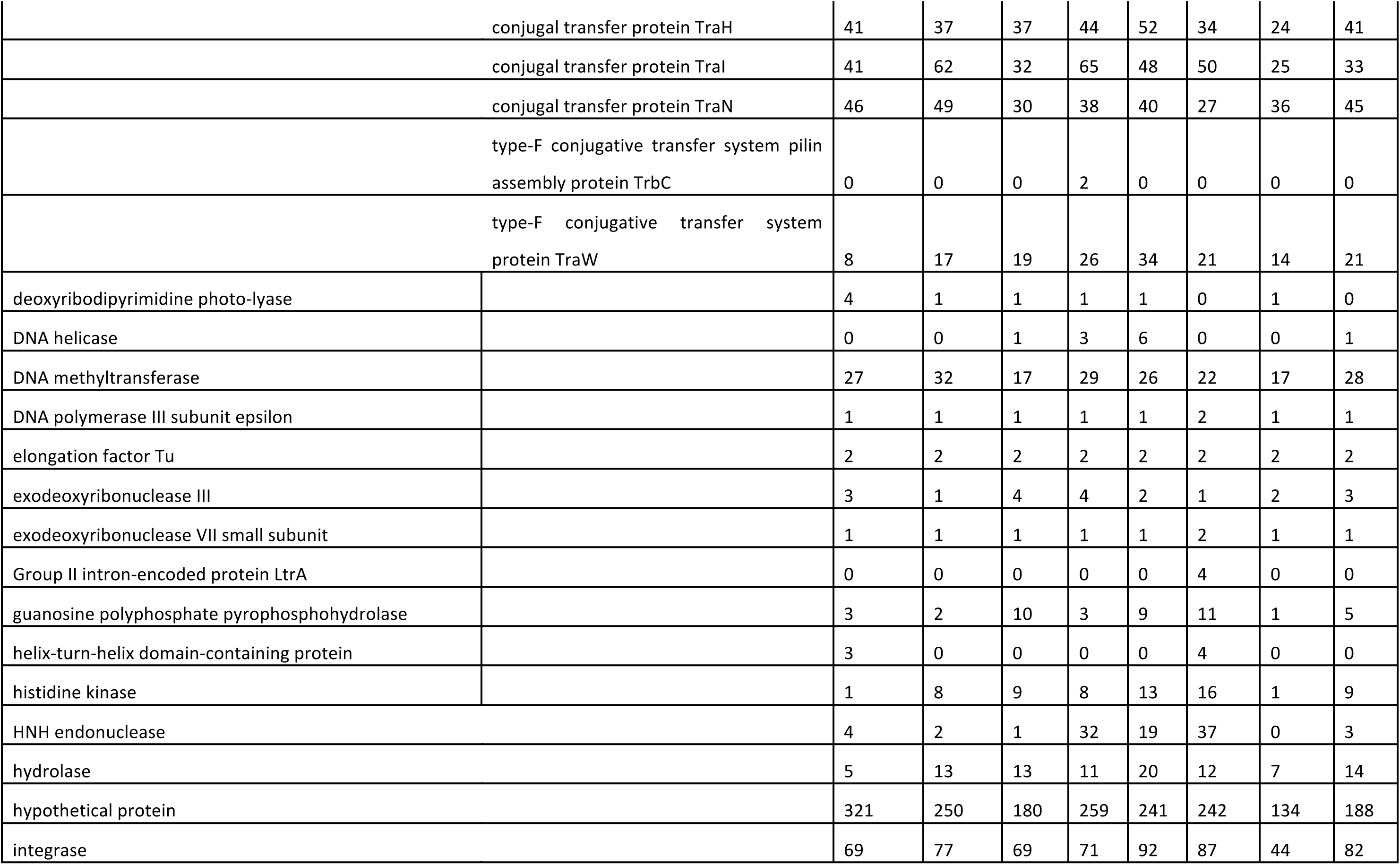
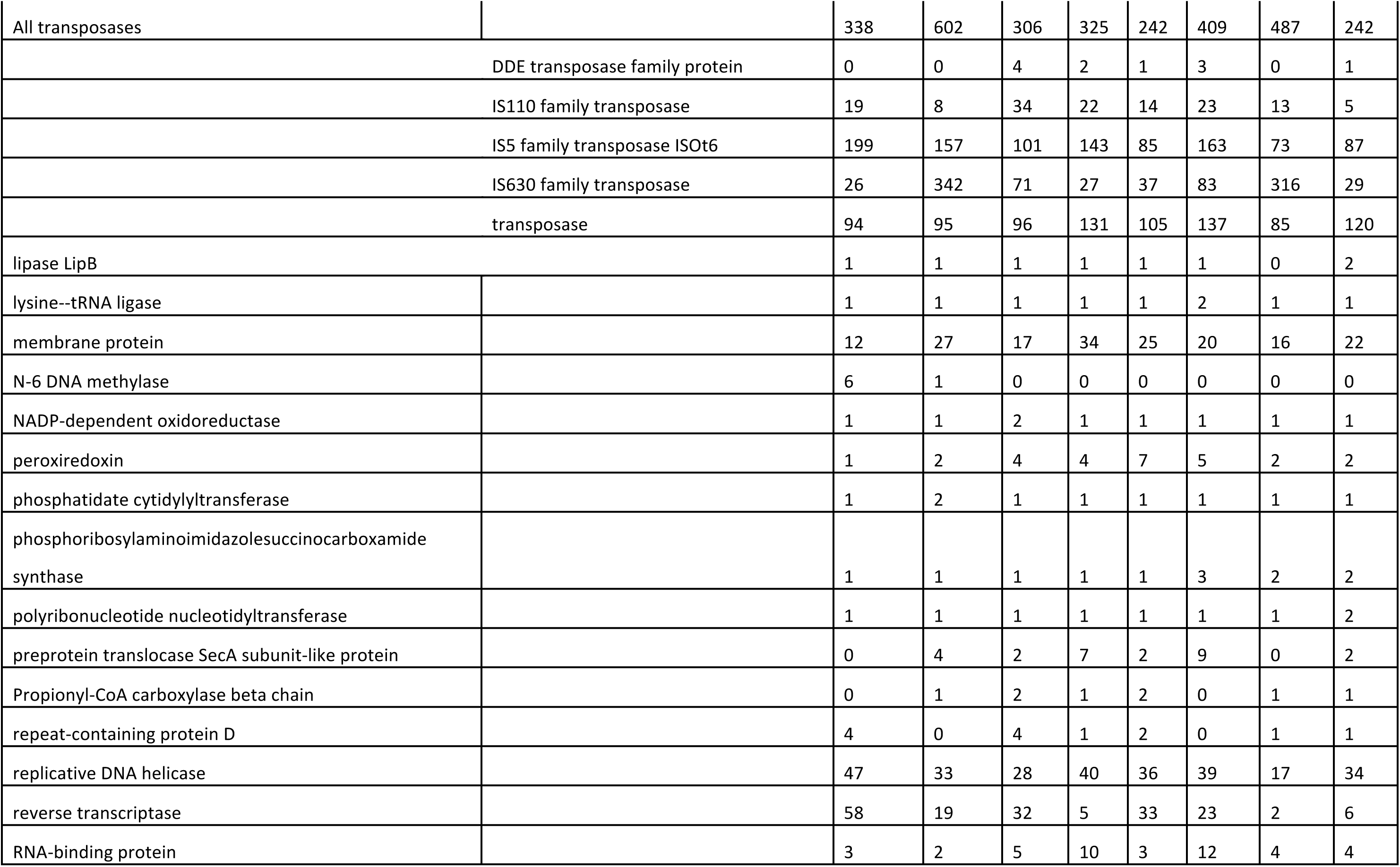
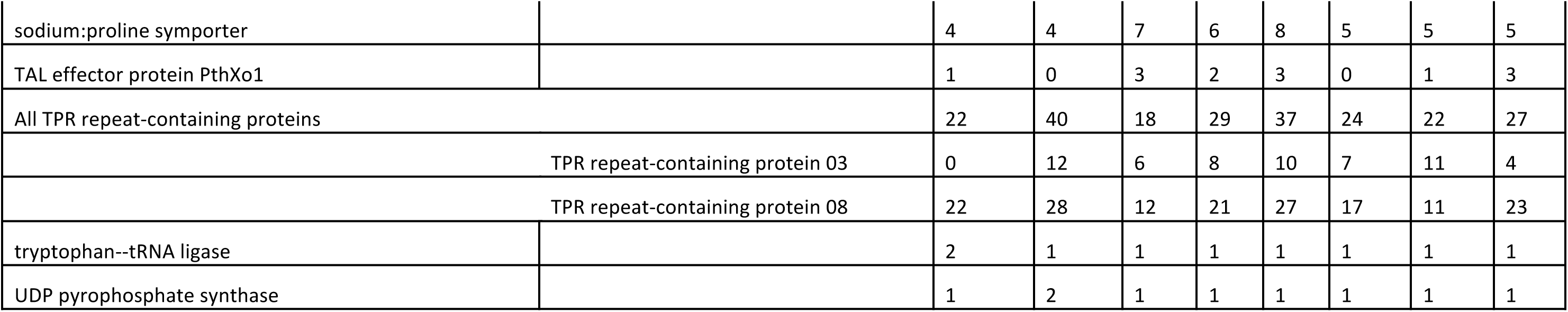
Repeat gene counts in each strain. Repeat genes were grouped by protein similarity and annotated with the product of the longest gene in the group where annotations differed.

**Table S7.** Pseudogenes and causes of pseudogenisation for each strain. The causes are not mutually exclusive, and may sum to greater than the total number of pseudogenes.

| Sample | Pseudogenes | Truncated 5' | Truncated 3' | Frameshift |
| --- | --- | --- | --- | --- |
| Boryong | 432 | 219 | 302 | 46 |
| Gilliam | 484 | 262 | 278 | 51 |
| Ikeda | 257 | 141 | 186 | 38 |
| Karp | 321 | 105 | 236 | 47 |
| Kato | 286 | 143 | 178 | 57 |
| TA686 | 453 | 200 | 307 | 50 |
| UT176 | 465 | 107 | 392 | 53 |
| UT76 | 319 | 149 | 203 | 52 |

**Table S8.** Robinson-Foulds distances between phylogenetic trees.

|  | 56kDa | 47kDa | MLST | Core genome |
| --- | --- | --- | --- | --- |
| 56kDa | - | 10 | 10 | 8 |
| 47kDa | 10 | - | 8 | 6 |
| MLST | 10 | 8 | - | 10 |
| Core genome | 8 | 6 | 10 | - |

## References

Abouelhoda, M.I., Kurtz, S., and Ohlebusch, E. (2004). Replacing suffix trees with enhanced suffix arrays. J. Discret. Algorithms 2, 53–86.

Altschul, S.F., Madden, T.L., Schäffer, A.A., Zhang, J., Zhang, Z., Miller, W., and Lipman, D.J. (1997). Gapped {BLAST} and {PSI-BLAST:} a new generation of protein database search programs. Nucleic Acids Res. 25, 3389–3402.

Arai, S., Tabara, K., Yamamoto, N., Fujita, H., Itagaki, A., Kon, M., Satoh, H., Araki, K., Tanaka-Taya, K., Takada, N., et al. (2013). Molecular phylogenetic analysis of Orientia tsutsugamushi based on the groES and groEL genes. Vector Borne Zoonotic Dis. 13, 825–829.

Blacksell, S.D., Luksameetanasan, R., Kalambaheti, T., Aukkanit, N., Paris, D.H., McGready, R., Nosten, F., Peacock, S.J., and Day, N.P.J. (2008). Genetic typing of the 56-kDa type-specific antigen gene of contemporary *Orientia tsutsugamushi* isolates causing human scrub typhus at two sites in north-eastern and western Thailand. FEMS Immunol. Med. Microbiol. 52, 335–342.

Bonell, A., Lubell, Y., Newton, P.N., Crump, J.A., and Paris, D.H. (2017). Estimating the burden of scrub typhus: A systematic review. PLoS Negl. Trop. Dis. 11, e0005838.

Cho, N.-H., Kim, H.-R., Lee, J.-H., Kim, S.-Y., Kim, J., Cha, S., Kim, S.-Y., Darby, A.C., Fuxelius, H.-H., Yin, J., et al. (2007). The Orientia tsutsugamushi genome reveals massive proliferation of conjugative type IV secretion system and host-cell interaction genes. Proc. Natl. Acad. Sci. U. S. A. 104, 7981–7986.

Cock, P.J.A., Antao, T., Chang, J.T., Chapman, B.A., Cox, C.J., Dalke, A., Friedberg, I., Hamelryck, T., Kauff, F., Wilczynski, B., et al. (2009). Biopython: freely available Python tools for computational molecular biology and bioinformatics. Bioinformatics 25, 1422–1423.

Coleman, R.E., Monkanna, T., Linthicum, K.J., Strickman, D.A., Frances, S.P., Tanskul, P., Kollars, T.M., Inlao, I., Watcharapichat, P., Khlaimanee, N., et al. (2003). Occurrence of Orientia tsutsugamushi in small mammals from Thailand. Am. J. Trop. Med. Hyg. 69, 519–524.

Cox, M.M., Keck, J.L., and Battista, J.R. (2010). Rising from the Ashes: DNA Repair in Deinococcus radiodurans. PLoS Genet. 6, e1000815.

Darby, A.C., Cho, N.-H., Fuxelius, H.-H., Westberg, J., and Andersson, S.G.E. (2007). Intracellular pathogens go extreme: genome evolution in the Rickettsiales. Trends Genet. 23, 511–520.

Dittrich, S., Rattanavong, S., Lee, S.J., Panyanivong, P., Craig, S.B., Tulsiani, S.M., Blacksell, S.D., Dance, D.A.B., Dubot-Pérès, A., Sengduangphachanh, A., et al. (2015). Orientia, rickettsia, and leptospira pathogens as causes of CNS infections in Laos: a prospective study. Lancet. Glob. Heal. 3, e104–12.

Duong, V., Blassdell, K., May, T.T.X., Sreyrath, L., Gavotte, L., Morand, S., Frutos, R., and Buchy, P. (2013). Diversity of Orientia tsutsugamushi clinical isolates in Cambodia reveals active selection and recombination process. Infect. Genet. Evol. 15, 25–34.

Enatsu, T., Urakami, H., and Tamura, A. (1999). Phylogenetic analysis of *Orientia tsutsugamushi* strains based on the sequence homologies of 56-kDa type-specific antigen genes. FEMS Microbiol. Lett. 180, 163–169.

Frutos, R., Viari, A., Ferraz, C., Morgat, A., Eychenié, S., Kandassamy, Y., Chantal, I., Bensaid, A., Coissac, E., Vachiery, N., et al. (2006). Comparative genomic analysis of three strains of Ehrlichia ruminantium reveals an active process of genome size plasticity. J. Bacteriol. 188, 2533–2542.

Fu, L., Niu, B., Zhu, Z., Wu, S., and Li, W. (2012). CD-HIT: accelerated for clustering the next-generation sequencing data. Bioinformatics 28, 3150–3152.

Giengkam, S., Blakes, A., Utsahajit, P., Chaemchuen, S., Atwal, S., Blacksell, S.D., Paris, D.H., Day, N.P.J., and Salje, J. (2015). Improved Quantification, Propagation, Purification and Storage of the Obligate Intracellular Human Pathogen Orientia tsutsugamushi. PLoS Negl. Trop. Dis. 9, e0004009.

Gillespie, J.J., Joardar, V., Williams, K.P., Driscoll, T., Hostetler, J.B., Nordberg, E., Shukla, M., Walenz, B., Hill, C.A., Nene, V.M., et al. (2012). A Rickettsia genome overrun by mobile genetic elements provides insight into the acquisition of genes characteristic of an obligate intracellular lifestyle. J. Bacteriol. 194, 376–394.

Izzard, L., Fuller, A., Blacksell, S.D., Paris, D.H., Richards, A.L., Aukkanit, N., Nguyen, C., Jiang, J., Fenwick, S., Day, N.P.J., et al. (2010). Isolation of a Novel *Orientia* Species (*O. chuto* sp. nov.) from a Patient Infected in Dubai. J. Clin. Microbiol. 48, 4404–4409.

Jiang, J., Paris, D.H., Blacksell, S.D., Aukkanit, N., Newton, P.N., Phetsouvanh, R., Izzard, L., Stenos, J., Graves, S.R., Day, N.P.J., et al. (2013). Diversity of the 47-kD HtrA nucleic acid and translated amino acid sequences from 17 recent human isolates of Orientia. Vector Borne Zoonotic Dis. 13, 367–375.

Jones, E., Oliphant, T., Peterson, P., and others {SciPy}: Open source scientific tools for {Python}.

Krzywinski, M., Schein, J., Birol, I., Connors, J., Gascoyne, R., Horsman, D., Jones, S.J., and Marra, M.A. (2009). Circos: An information aesthetic for comparative genomics. Genome Res. 19, 1639–1645.

Leimbach, A. (2016). bac-genomics-scripts: Bovine E. coli mastitis comparative genomics edition.

Li, W., and Godzik, A. (2006). Cd-hit: a fast program for clustering and comparing large sets of protein or nucleotide sequences. Bioinformatics 22, 1658–1659.

Liao, H.-M., Chao, C.-C., Lei, H., Li, B., Tsai, S., Hung, G.-C., Ching, W.-M., and Lo, S.-C. (2017). Intraspecies comparative genomics of three strains of Orientia tsutsugamushi with different antibiotic sensitivity. Genomics Data 12, 84–88.

Lu, H.-Y., Tsai, K.-H., Yu, S.-K., Cheng, C.-H., Yang, J.-S., Su, C.-L., Hu, H.-C., Wang, H.-C., Huang, J.-H., and Shu, P.-Y. (2010). Phylogenetic analysis of 56-kDa type-specific antigen gene of Orientia tsutsugamushi isolates in Taiwan. Am. J. Trop. Med. Hyg. 83, 658–663.

Luce-Fedrow, A., Lehman, M., Kelly, D., Mullins, K., Maina, A., Stewart, R., Ge, H., John, H., Jiang, J., and Richards, A. (2018). A Review of Scrub Typhus (Orientia tsutsugamushi and Related Organisms): Then, Now, and Tomorrow. Trop. Med. Infect. Dis. 3, 8.

Lunter, G., and Goodson, M. (2010). Stampy: A statistical algorithm for sensitive and fast mapping of Illumina sequence reads. Genome Res.

Marchler-Bauer, A., Panchenko, A.R., Shoemaker, B.A., Thiessen, P.A., Geer, L.Y., and Bryant, S.H. (2002). CDD: a database of conserved domain alignments with links to domain three-dimensional structure. Nucleic Acids Res. 30, 281–283.

McGready, R., Blacksell, S.D., Luksameetanasan, R., Wuthiekanun, V., Jedsadapanpong, W., Day, N.P.J., and Nosten, F. (2010). First report of an Orientia tsutsugamushi type TA716-related scrub typhus infection in Thailand. Vector Borne Zoonotic Dis. 10, 191–193.

Merhej, V., and Raoult, D. (2011). Rickettsial evolution in the light of comparative genomics. Biol. Rev. Camb. Philos. Soc. 86, 379–405.

Moran, N.A. (1996). Accelerated evolution and Muller’s rachet in endosymbiotic bacteria. Proc. Natl. Acad. Sci. U. S. A. 93, 2873–2878.

Nakayama, K., Yamashita, A., Kurokawa, K., Morimoto, T., Ogawa, M., Fukuhara, M., Urakami, H., Ohnishi, M., Uchiyama, I., Ogura, Y., et al. (2008). The Whole-genome sequencing of the obligate intracellular bacterium Orientia tsutsugamushi revealed massive gene amplification during reductive genome evolution. DNA Res. 15, 185–199.

Nakayama, K., Kurokawa, K., Fukuhara, M., Urakami, H., Yamamoto, S., Yamazaki, K., Ogura, Y., Ooka, T., and Hayashi, T. (2010). Genome comparison and phylogenetic analysis of Orientia tsutsugamushi strains. DNA Res. 17, 281–291.

Page, A.J., Cummins, C.A., Hunt, M., Wong, V.K., Reuter, S., Holden, M.T.G., Fookes, M., Falush, D., Keane, J.A., and Parkhill, J. (2015). Roary: rapid large-scale prokaryote pan genome analysis. Bioinformatics 31, 3691–3693.

Paradis, E., Claude, J., and Strimmer, K. (2004). A{PE}: analyses of phylogenetics and evolution in {R} language. Bioinformatics 20, 289–290.

Paris, D.H., Aukkanit, N., Jenjaroen, K., Blacksell, S.D., and Day, N.P.J. (2009). A highly sensitive quantitative real-time PCR assay based on the groEL gene of contemporary Thai strains of Orientia tsutsugamushi. Clin. Microbiol. Infect. 15, 488–495.

Paris, D.H., Phetsouvanh, R., Tanganuchitcharnchai, A., Jones, M., Jenjaroen, K., Vongsouvath, M., Ferguson, D.P.J., Blacksell, S.D., Newton, P.N., Day, N.P.J., et al. (2012). Orientia tsutsugamushi in human scrub typhus eschars shows tropism for dendritic cells and monocytes rather than endothelium. PLoS Negl. Trop. Dis. 6, e1466.

Phetsouvanh, R., Sonthayanon, P., Pukrittayakamee, S., Paris, D.H., Newton, P.N., Feil, E.J., Day, N.P.J., Kurup, A., Issac, A., Loh, J., et al. (2015). The Diversity and Geographical Structure of Orientia tsutsugamushi Strains from Scrub Typhus Patients in Laos. PLoS Negl. Trop. Dis. 9, e0004024.

Pritchard, L., White, J.A., Birch, P.R.J., and Toth, I.K. (2006). GenomeDiagram: a python package for the visualization of large-scale genomic data. Bioinformatics 22, 616–617.

R Core Team (2014). R: A Language and Environment for Statistical Computing.

Revell, L.J. (2012). phytools: An R package for phylogenetic comparative biology (and other things). Methods Ecol. Evol. 3, 217–223.

Rights, F.L., and Smadel, J.E. (1948). Studies on scrub typhus; tsutsugamushi disease; heterogenicity of strains of R. tsutsugamushi as demonstrated by cross-vaccination studies. J. Exp. Med. 87, 339–351.

Schliep, K.P. (2011). phangorn: phylogenetic analysis in R. Bioinformatics 27, 592–593.

Seemann, T. (2014). Prokka: rapid prokaryotic genome annotation. Bioinformatics 30, 2068–2069.

Sievers, F., Wilm, A., Dineen, D., Gibson, T.J., Karplus, K., Li, W., Lopez, R., McWilliam, H., Remmert, M., Söding, J., et al. (2011). Fast, scalable generation of high-quality protein multiple sequence alignments using Clustal Omega. Mol. Syst. Biol. 7.

Sonthayanon, P., Peacock, S.J., Chierakul, W., Wuthiekanun, V., Blacksell, S.D., Holden, M.T.G., Bentley, S.D., Feil, E.J., and Day, N.P.J. (2010). High rates of homologous recombination in the mite endosymbiont and opportunistic human pathogen Orientia tsutsugamushi. PLoS Negl. Trop. Dis. 4, e752.

Stamatakis, A. (2014). RAxML version 8: a tool for phylogenetic analysis and post-analysis of large phylogenies. Bioinformatics 30, 1312–1313.

Walker, B.J., Abeel, T., Shea, T., Priest, M., Abouelliel, A., Sakthikumar, S., Cuomo, C.A., Zeng, Q., Wortman, J., Young, S.K., et al. (2014). Pilon: An Integrated Tool for Comprehensive Microbial Variant Detection and Genome Assembly Improvement. PLoS One 9, e112963.

Weitzel, T., Dittrich, S., López, J., Phuklia, W., Martinez-Valdebenito, C., Velásquez, K., Blacksell, S.D., Paris, D.H., and Abarca, K. (2016). Endemic Scrub Typhus in South America. N. Engl. J. Med. 375, 954–961.

Wiens, G.D., Rockey, D.D., Wu, Z., Chang, J., Levy, R., Crane, S., Chen, D.S., Capri, G.R., Burnett, J.R., Sudheesh, P.S., et al. (2008). Genome sequence of the fish pathogen Renibacterium salmoninarum suggests reductive evolution away from an environmental Arthrobacter ancestor. J. Bacteriol. 190, 6970–6982.

Wongprompitak, P., Duong, V., Anukool, W., Sreyrath, L., Mai, T.T.X., Gavotte, L., Moulia, C., Cornillot, E., Ekpo, P., Suputtamongkol, Y., et al. (2015). Orientia tsutsugamushi, agent of scrub typhus, displays a single metapopulation with maintenance of ancestral haplotypes throughout continental South East Asia. Infect. Genet. Evol. 31, 1–8.

Wu, M., Sun, L. V, Vamathevan, J., Riegler, M., Deboy, R., Brownlie, J.C., McGraw, E.A., Martin, W., Esser, C., Ahmadinejad, N., et al. (2004). Phylogenomics of the Reproductive Parasite Wolbachia pipientis wMel: A Streamlined Genome Overrun by Mobile Genetic Elements. PLoS Biol. 2, e69.

